# Microbiota-induced plastic T cells enhance immune control of antigen-sharing tumors

**DOI:** 10.1101/2024.08.12.607605

**Authors:** Tariq A. Najar, Yuan Hao, Yuhan Hao, Gabriela Romero-Meza, Alexandra Dolynuk, Dan R. Littman

## Abstract

Therapies that harness the immune system to target and eliminate tumor cells have revolutionized cancer care. Immune checkpoint blockade (ICB), which boosts the anti-tumor immune response by inhibiting negative regulators of T cell activation^1–3^, is remarkably successful in a subset of cancer patients, yet a significant proportion do not respond to treatment, emphasizing the need to understand factors influencing the therapeutic efficacy of ICB^4–9^. The gut microbiota, consisting of trillions of microorganisms residing in the gastrointestinal tract, has emerged as a critical determinant of immune function and response to cancer immunotherapy, with multiple studies demonstrating association of microbiota composition with clinical response^10–16^. However, a mechanistic understanding of how gut commensal bacteria influence the efficacy of ICB remains elusive. Here we utilized a gut commensal microorganism, segmented filamentous bacteria (SFB), which induces an antigen-specific Th17 cell effector program^17^, to investigate how colonization with it affects the efficacy of ICB in restraining distal growth of tumors sharing antigen with SFB. We find that anti-PD-1 treatment effectively inhibits the growth of implanted SFB antigen-expressing melanoma only if mice are colonized with SFB. Through T cell receptor clonal lineage tracing, fate mapping, and peptide-MHC tetramer staining, we identify tumor-associated SFB-specific Th1-like cells derived from the homeostatic Th17 cells induced by SFB colonization in the small intestine lamina propria. These gut-educated ex-Th17 cells produce high levels of the pro-inflammatory cytokines IFN-γ and TNF-α, and promote expansion and effector functions of CD8^+^ tumor-infiltrating cytotoxic lymphocytes, thereby controlling tumor growth. A better understanding of how distinct intestinal commensal microbes can promote T cell plasticity-dependent responses against antigen-sharing tumors may allow for the design of novel cancer immunotherapeutic strategies.

## MAIN

Although specific bacterial species have been associated with favorable ICB treatment responses in cancer patients^12,13,18–22^, there is little understanding of mechanisms by which the intestinal microbiota composition influences anti-tumor immune responses. Products of the microbiota, including metabolites and ligands for innate immune receptors, may contribute to enhancing the functions of antigen-presenting cells, thus lowering the threshold for activation of tumor-reactive T cells. Alternatively, T cells specific for antigens shared by the microbiota and tumor cells may become activated in the setting of ICB, thus enhancing anti-tumor responses. Because the microbiome encompasses a vast number of antigens with the potential to be presented to host T cells, there is a strong possibility that some bacteria with access to the host immune system elicit T cells with receptor cross-reactivity for tumor antigens, resulting in control of tumor growth. Despite correlative data^23^, the causality of such antigenic mimicry has not yet been definitively demonstrated in cancer patients. It is possible, however, to test this premise, particularly the relationship of microbe-specific T cells and intratumoral T cells, in animal models. An immunization model with skin-associated *S. epidermidis* engineered to express antigens shared with implanted tumors was shown to elicit effective anti-tumor responses^24^, but the ability of gut commensals that elicit stereotyped T cell responses to program anti-tumor immunity has not been explored. Here, we studied how a small intestine-resident commensal microbe, SFB, that induces a regulatory-like Th17 cell response that enhances intestinal barrier integrity^25,26^, influences efficacy of ICB in controlling growth of distal tumors that share antigen with the bacterium. We found that tumor-specific Th17 cells primed by SFB in the gut infiltrate the tumor as trans-differentiated Th1-like cells following ICB. These cells promote recruitment and maturation of CD8^+^ effector T cells that contribute critically to anti-tumor immunity. Our results suggest that defined constituents of the intestinal microbiota can be harnessed to elicit desired effector T cell programs that restrain tumor growth.

## RESULTS

### SFB promotes ICB-mediated tumor control

To explore how commensal microbiota can contribute to immune-mediated tumor control, we developed a synthetic neoantigen mimicry tumor model in mice by expressing an antigen from gut-colonizing SFB in B16-F10 melanoma cells (**Figure 1a**). We chose SFB because it is a non-pathogenic commensal microbe that elicits a well-characterized immune response with local beneficial functions as well as systemic effects^17,27,28^. We expressed a fragment of the bacterial protein SFB-3340 (**Figure 1b**) that elicits in SFB-colonized mice a dominant CD4^+^ T cell response that can be monitored using peptide-MHCII tetramers and T cell antigen receptor transgenic mice. Extracts from tumor cells expressing the SFB protein (B16-3340), but not from cells transduced with an empty vector (B16-EV), activated the TCR transgenic T cells *ex vivo* (**Figure 1c**).

**Figure 1.**
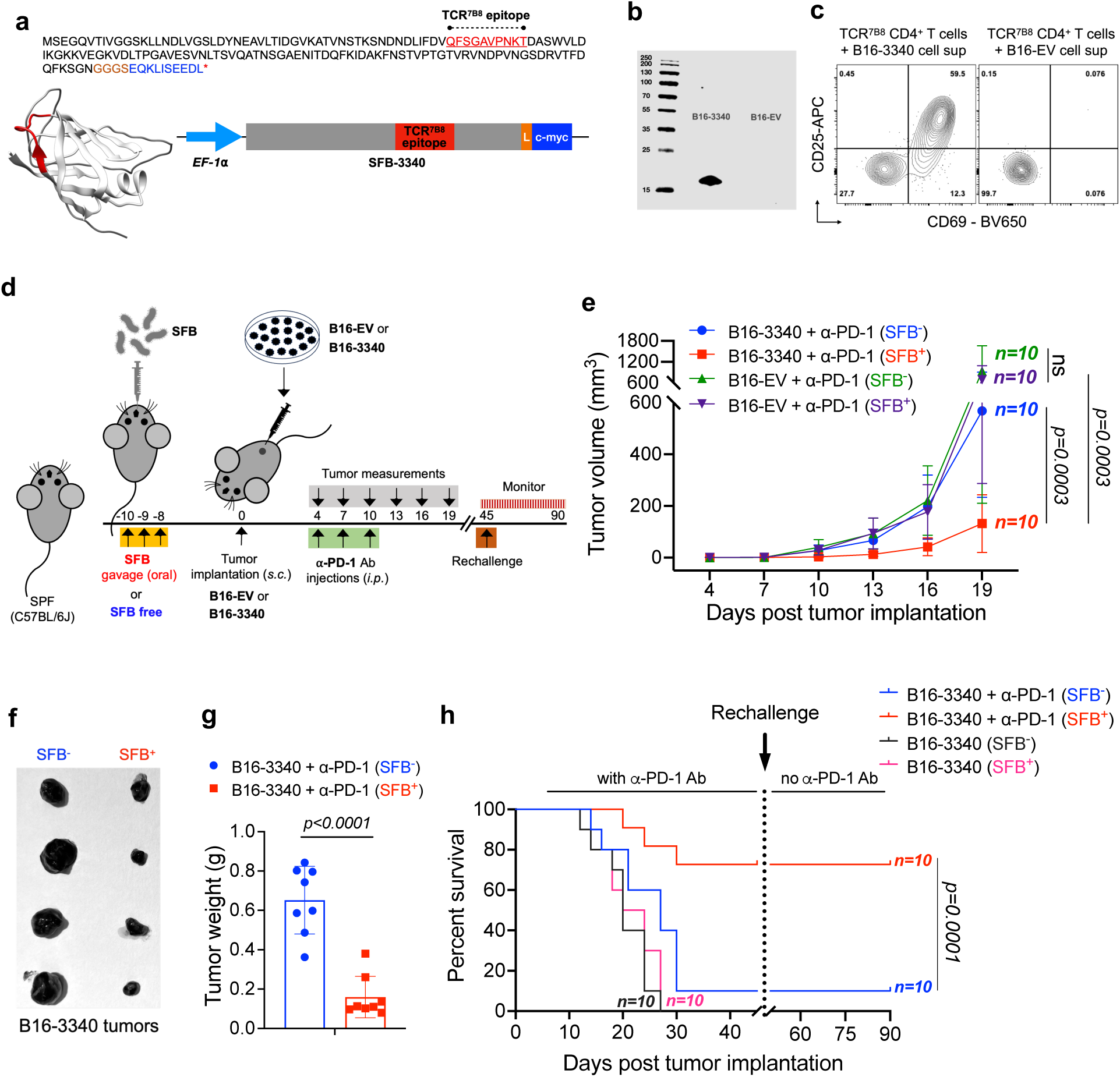
Development of a synthetic microbiota-based tumor antigen mimicry model to evaluate the response to anti-PD1 antibody treatment. **(a)** Amino acid sequence of a segment of an independently folding domain of the SFB-3340 protein, containing the CD4^+^ T cell (TCR^7B8^) epitope (red). The codon optimized gene was fused in frame with a EF1-α promoter on the 5’-end and c-myc tag (blue) at the 3’-end by a flexible linker (L). **(b)** Western blot confirming the expression of the SFB-3340 protein fragment in transfected B16-F10 (B16-3340) cells, detected by anti c-myc antibody staining. The control B16-F10 cell line was transfected with empty vector (B16-EV). (c) *Ex-vivo* activation of *naïve* SFB-specific CD4^+^ T cells from TCR^7B8^ transgenic mice following incubation with either cell supernatant from lysed B16-3340 or B16-EV cells and syngeneic splenocytes. Activation markers CD69 and CD25 were detected at 24h. **(d)** Experimental strategy for testing the artificial mimicry model in SPF C57BL/J mice, comparing groups that were either colonized with SFB or kept SFB-free. **(e)** Caliper measurements are shown as growth curves of B16-3340 or B16-EV implanted tumors. Mice in each group were either colonized with SFB (n=10) or kept SFB-free (n=10) and subcutaneously implanted with either B16-3340 or B16-EV cells. All mice received 3 injections of anti-PD-1 antibody (250 µg/mouse *i.p.* on days 4, 7 and 10 post tumor implantation). Statistical significance was calculated using two-way ANOVA and Sidak’s multiple comparisons. **(f)** Comparison of excised B16-3340 tumor tissues from control (SFB-free) and SFB-colonized mice at day 14 post tumor implantation. **(g)** Quantification of excised tumor tissue weights from SFB^-^ and SFB^+^ mice at day 14 post tumor implantation (n=8). Statistical significance was determined using unpaired two-sided Mann-Whitney t-test. Error bars denote mean ± SD. *P* values are indicated on the corresponding graphs. **(h)** Kaplan-Meier survival curves comparing SFB-colonized and control mice bearing B16-3340 tumors, with and without anti-PD-1 treatment. Following the initial challenge, surviving mice were re-challenged with the same tumor cells, and subsequent survival was monitored without anti-PD-1 antibody treatment. *P* value generated by the log-rank (Mantel-Cox) test.

Next, we examined the effect of SFB colonization on tumor growth in specific pathogen-free (SPF) mice (Jackson Laboratories) bearing subcutaneously implanted B16-3340 and B16-EV tumors, with (**Figure 1d**) or without (**Extended Data Fig. 1a**) anti-PD-1 antibody treatment. In the absence of PD-1 blockade, there was no significant difference in tumor growth between mice implanted with either B16-3340 or B16-EV, regardless of SFB colonization status (SFB^+^ or SFB^-^) (**Extended Data Fig. 1b, c**). However, when animals were treated with anti-PD-1 antibody, the growth of B16-3340 tumors was significantly reduced in SFB^+^ mice compared to SFB^-^ mice. There was no notable difference in the growth of control B16-EV tumors between SFB^+^ and SFB^-^ mice receiving anti-PD-1 antibody treatment (**Figure 1e-g and Extended Data Fig. 1d).** The combination of SFB colonization and anti-PD-1 treatment of B16-3340 tumors also resulted in significantly higher survival rates compared to the other groups (**Figure 1h**). Furthermore, when these treated, now tumor-free, mice were subsequently re-challenged with B16-3340 cells, they completely rejected the tumor without requiring additional anti-PD-1 treatment, suggesting that a T cell memory response elicited by earlier SFB colonization, in combination with checkpoint blockade, was sufficient to restrict tumor growth. The enhanced response to anti-PD-1 treatment observed in SFB-colonized mice, compared to SFB-free mice, prompted further investigation into the mechanisms by which gut microbiota, specifically SFB, influences anti-tumor immune responses and augments the efficacy of ICB in treated animals.

### SFB alters tumor T cell composition and phenotype

First, we examined the composition and effector functions of T cell in tumors (B16-3340 and B16-EV) from SFB^+^ and SFB^-^ mice, all subjected to anti-PD-1 antibody treatment (**Figure 2a**). Within B16-3340 tumors in SFB-colonized mice, there was a significant increase in the CD8^+^ to regulatory T (Treg) cell ratio compared to either SFB^-^ mice with B16-3340 tumors or SFB^+^ mice with B16-EV tumors (**Figure 2b**). In contrast, there was no difference in the CD8^+^ to Treg ratio in the small intestine lamina propria (SILP) of SFB^+^ and SFB^-^ mice treated with anti-PD-1 antibody (**Extended Data Fig. 2a**). The combination of SFB colonization and anti-PD-1 treatment significantly enhanced CD8^+^ T cell effector functions, with increased frequencies of IFN-γ^+^, TNF-α^+^IFN-γ^+^, and granzyme B^+^TNF-α^+^ CD8^+^ T cells in B16-3340 tumors. Similar results were previously demonstrated in the microbiota-mediated response to PD-1 blockade^29^. In contrast, CD8^+^ tumor-infiltrating lymphocytes (TILs) from either B16-3340 tumors in SFB^-^ mice or B16-EV tumors in SFB^+^ mice failed to produce these effector molecules (**Figure 2c and Extended Data Fig. 2b**).

**Figure 2:**
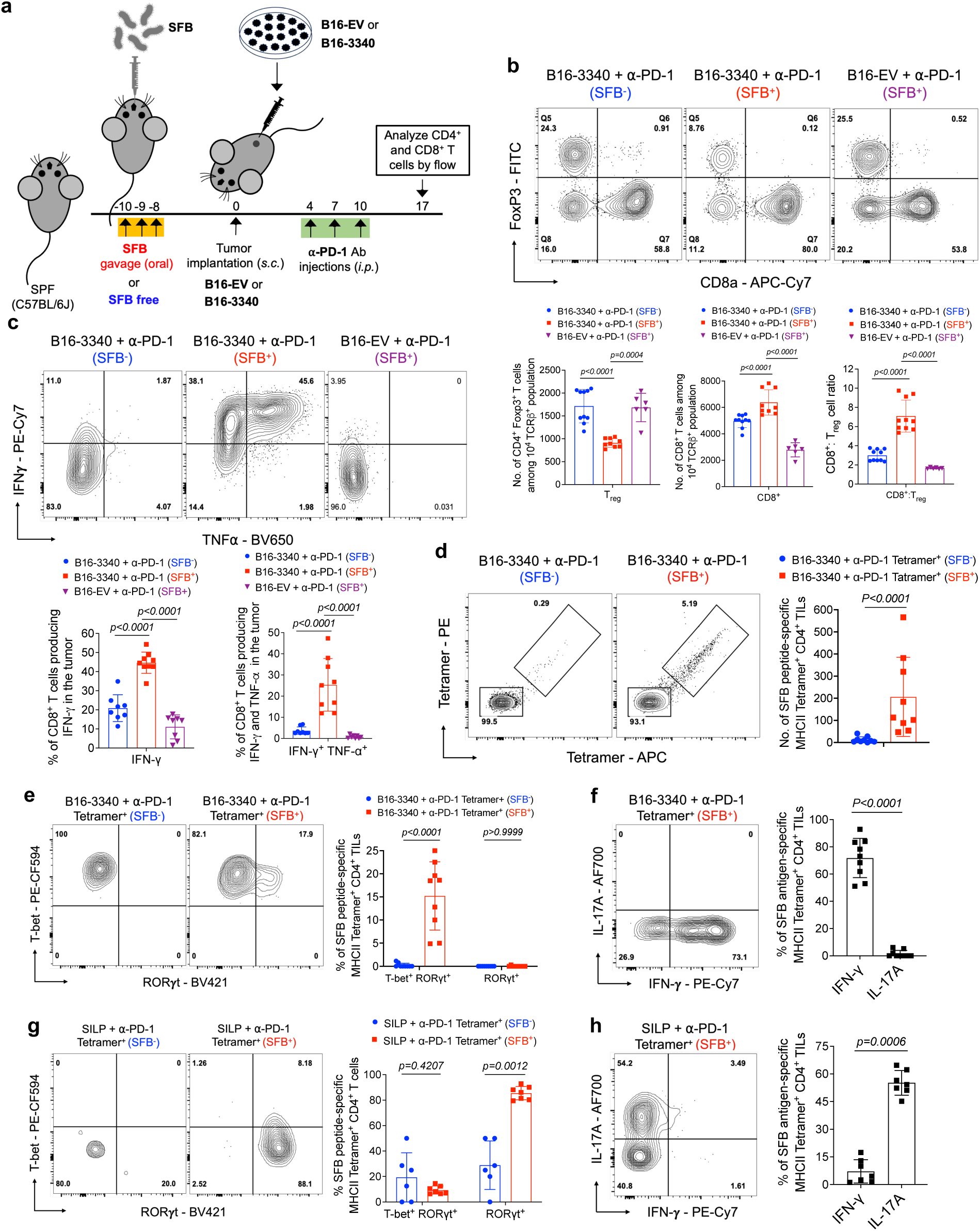
SFB colonization modulates effector functions of both CD8^+^ and CD4^+^ tumor infiltrating T cells. **(a)** Schematic representation of the synthetic mimicry model to examine the effects of gut colonizing commensal microbe (SFB) on the distal tumor microenvironment. **(b)** The top panel displays representative flow cytometry plots of CD8^+^ T cells and regulatory T cells (Tregs; CD4^+^ Foxp3^+^) in B16-3340 tumors from SFB-free (SFB^-^) and SFB-colonized (SFB^+^) mice (n=10 per group), as well as in B16-EV tumors from SFB-colonized mice (n=6). The bottom panel quantifies the cell number of CD8^+^ T cells and Tregs, as well as the CD8 to Treg cell ratio across these groups. **(c)** The top panel displays representative flow cytometry plots showing the expression of effector gene products (TNF-α^+^ IFN-γ^+^) in CD8^+^ TILs isolated from B16-3340 tumors in SFB^-^ and SFB^+^ mice, and B16-EV tumors in SFB^+^ mice. The bottom panel quantifies the percentage of CD8^+^ TILs expressing both TNF-α and IFN-γ (double-positive cells) across the different groups (n=8 to 9 per group). **(d)** SFB peptide-specific MHCII tetramer staining of CD4^+^ TILs isolated from B16-3340 implanted tumors of either SFB^-^ or SFB^+^ mice treated with anti-PD-1 antibody (n=9 per group). (**e**) Representative flow cytometry plots of transcription factor, RORγt and T-bet expression profiles of Tetramer^+^ CD4^+^ T cells isolated from the B16-3340 tumors of SFB^-^ and SFB^+^ mice (left panel) and quantification of transcription factor expression across the two groups (right panel, n=8 to 9 per group). (**f**) Expression of effector gene products, IFN-γ and IL-17A, in Tetramer^+^ CD4^+^ T cells from B16-3340 tumors in SFB^+^ mice (left panel) and quantification (right panel, n=9 per group). (**g**) Representative flow cytometry plots displaying the expression of transcription factors, RORγt and T-bet in Tetramer^+^ CD4^+^ T cells isolated from the SILP of SFB^-^ and SFB^+^ mice (left panel). The right panel illustrates the quantification of transcription factor expression across the two groups (n=6 to 7 per group). (**h**) Analysis of effector gene products, IFN-γ and IL-17A, expression in Tetramer^+^ CD4^+^ T cells from SILP in SFB^+^ mice (left panel) and quantification (right panel, n=7 per group). In each experiment, three doses of anti-PD-1 antibody were administered to the animals in both the SFB^-^ and SFB^+^ groups. For cytokine expression analysis, cells were activated *ex vivo* by PMA/Ionomycin for 3 hours at 37°C. Statistical significance was determined using unpaired two-sided Mann-Whitney t-test. Error bars denote mean ± SD. *P* values are indicated on the corresponding bar graphs.

Next, because SFB colonization induces antigen-specific Th17 cells in the ileal lamina propria, we compared the CD4^+^ T cell phenotypes in the remotely-located B16-3340 tumors. First, using a panel of antibodies specific for TCR Vβs, we found a greater proportion of Vβ14^+^ CD4^+^ T cells in tumors from SFB^+^ compared to SFB^-^ mice (**Extended Data Fig. 2c**). This bias is consistent with the known preferential interaction of this subset of TCRs with immunodominant SFB peptides (**Extended Data Fig. 2d**)^30^. Second, using SFB-3340 peptide-loaded MHC II tetramers revealed that colonization with SFB caused increased infiltration of SFB-3340-specific CD4^+^ T cells into B16-3340 tumors **(Figure 2d)**. Remarkably, these tetramer positive (Tetramer^+^) CD4^+^ T cells displayed an IFN-γ producing Th1-like phenotype, unlike SILP Tetramer^+^ T cells that, as expected, were IL-17A producing Th17 cells (**Figure 2e-h and Extended Data Fig. 2e-g**). Tumor-associated Tetramer^+^ T cells expressed the transcription factor T-bet and produced IFN-γ in both SFB^+^ and SFB^-^ mice, although a small proportion of cells from SFB-colonized mice additionally expressed RORγt (**Figure 2e, f and Extended Data Fig. 2f, g)**. In contrast, the majority of tetramer negative (Tetramer^−^) CD4^+^ T cells isolated from B16-3340 and B16-EV tumors, irrespective of SFB colonization, were regulatory-like T cells, expressing both T-bet and Foxp3 (**Extended Data Fig. 2h, i**), a phenotype associated with strong suppression of anti-tumor immune responses^31^.

In ELISpot assays, IFN-γ producing CD4^+^ TILs were significantly enriched in B16-3340 tumors from SFB^+^ mice compared to those from SFB^-^ mice (**Extended Data Fig. 3a**), and, in accord, Tetramer^+^ CD4^+^ TILs from those tumors exhibited a robust Th1 cytokine (IFN-γ and TNF-α) response following *ex vivo* stimulation (**Extended Data Fig. 3b**). Furthermore, while the frequency of both T-bet^+^Foxp3^-^ and T-bet^+^Foxp3^+^ cells among total CD4^+^ Tetramer^−^ T cell population was comparable, B16-3340 tumors in SFB^+^ animals had a significantly higher percentage of CD4^+^ Tetramer^−^ T cells that produced moderate amounts of IFN-γ and TNF-α following *ex vivo* stimulation with PMA/Ionomycin than either B16-3340 tumors in the SFB^-^ group or B16-EV tumors of SFB^+^ mice (**Extended Data Fig. 3c, d**). This suggests that SFB-specific pro-inflammatory CD4^+^ T cells in the tumor may contribute to altering the tumor microenvironment, making it more responsive to anti-PD-1 therapy.

Next, given that effective tumor growth control requires a synergistic effect involving both CD4^+^ and CD8^+^ T cells working in conjunction with ICB^32–34^, we aimed to investigate whether the combination of SFB colonization-induced CD4^+^ and tumor-infiltrating CD8^+^ T cells, together with anti-PD-1 treatment, is essential for controlling the growth of SFB-3340-expressing tumors. Following depletion of either CD4^+^ (**Extended Data Fig. 4a**) or CD8^+^ (**Extended Data Fig. 5a**) T cells in SFB-colonized, B16-3340 tumor-bearing mice, there was a significant loss in the efficacy of anti-PD-1 treatment (**Extended Data Fig. 4b** and **Extended Data Fig. 5b**). We then examined how depletion of CD4^+^ T cells impacts the tumor-controlling potential of CD8^+^ TILs. CD8^+^ T cells were isolated from B16-3340 tumors in SFB-colonized mice that had been treated with anti-PD-1 antibody with and without depletion of CD4^+^ T cells. The production of IFN-γ, TNF-α and granzyme B by the CD8^+^ TILs was significantly reduced in B16-3340 tumors of CD4 depleted mice compared to that of cells from control (non-CD4 depleted) mice, consistent with a critical role for microbiota-dependent CD4^+^ T cell help in the maturation and function of cytotoxic T cells within the tumor microenvironment (**Extended Data Fig. 4c-e**).

Similarly, we examined the effector functions of tumor-infiltrating CD4^+^ T cells in CD8^+^ T cell-depleted mice. The production of IFN-γ and TNF-α by the CD4^+^ TILs isolated from the B16-3340 tumors of CD8^+^ T cell-depleted mice was also reduced, compared to that of cells from control (non-CD8 depleted) mice, although the reduction was not as pronounced as that observed in CD8^+^ TILs from CD4^+^ T cell-depleted mice (**Extended Data Fig. 5c, d**).

Overall, our results demonstrate that SFB colonization, when combined with anti-PD-1 treatment, significantly enhances the immune response against tumors with shared antigen. Specifically, we observed an increased CD8^+^ to Treg cell ratio, heightened effector functions of CD8^+^ TILs, and the presence of SFB-3340-specific CD4^+^ T cells with pro-inflammatory properties in the tumor microenvironment. This indicates a robust synergy between microbiota-induced T cell responses and ICB. Notably, the depletion of either CD4^+^ or CD8^+^ T cells significantly compromised the efficacy of anti-PD-1 therapy, highlighting the critical role of SFB-induced CD4^+^ T cells in facilitating the maturation and function of cytotoxic CD8^+^ T cells within the tumor.

### Clonal T cells in gut and tumors post-SFB

To examine the relationship of intestinal and tumor-infiltrating T cells in SFB-colonized and non-colonized mice with B16-3340 tumors, we performed single-cell RNA sequencing and TCR repertoire analysis (scRNA-seq + scTCR-seq) with sorted CD4^+^ T cells from these tissues (**Figure 3a**). Unsupervised clustering of the scRNA-seq data revealed transcriptionally distinct subsets, with 9 and 10 clusters in SILP and B16-3340 tumor, respectively, each expressing known markers of different CD4^+^ T cell subsets (**Extended Data Fig. 6a**). As anticipated, an enrichment of the IL-17A^+^ Th17 subset (cluster 2) was observed within the SILP of SFB^+^ mice compared to SFB-free mice (**Figure 3b**). Similarly, enrichment of the IFN-γ^+^ Th1-like subset (cluster 1) was observed only in tumors from SFB colonized mice (**Figure 3c**), highlighting the distinct transcriptional profiles and subset distributions of CD4^+^ T cells in the intestinal and tumor environments of SFB-colonized mice, which likely play a crucial role in mediating the observed anti-tumor immune response in SFB-colonized mice.

**Figure 3:**
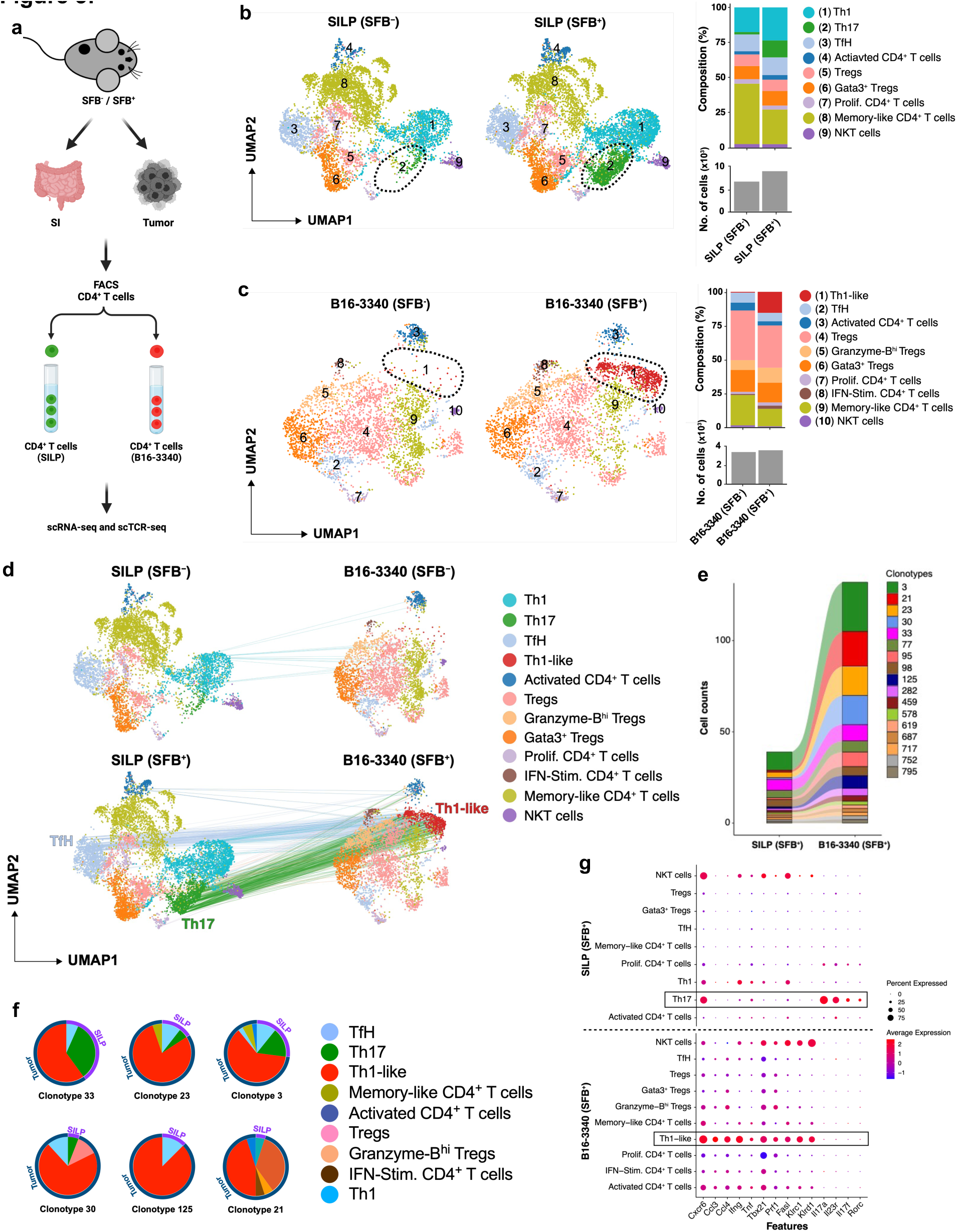
Shared T cell receptor (TCR) clonotypes in small intestine and tumors of SFB colonized and SFB-free mice. **(a)** Strategy for scRNA and scTCRseq of CD4^+^ T cells from B16-3340 tumor and SILP of anti-PD1 treated mice with and without SFB colonization. (**b**) UMAP visualization of sorted CD4^+^ T cell subsets from the SILP of SFB^-^ (left) and SFB^+^ (middle) mice. Right, CD4^+^ T cell subset distribution (top) and total numbers of T cells profiled (bottom). (**c**) UMAP visualization of sorted CD4^+^ T cell subsets from tumors of SFB^-^ (left) and SFB^+^ (middle) mice. The distribution and total numbers of CD4^+^ T cell subsets is shown on the right. **(d)** UMAPs illustrating the clonal connectivity between CD4^+^ T cells within the SILP and tumor tissues, providing a qualitative representation of their interrelationship. **(e)** Alluvial graph depicting the clonal connectivity of CD4^+^ T cell clonotypes between the SILP and tumor tissues in mice colonized with SFB. Clonality was determined based on identity of CDR3 nucleotide sequences for both TCR α and β paired chains. Each block in the bar diagram represents cell counts within a distinct CD4^+^ T cell clonotype, with branches in the graph illustrating the shared clonotypes between SILP and tumor compartments. (**f**) Clonal expansion with phenotypic switching (represented by color) within the tumor for select shared clonotypes originating from the gut. (**g)** Dot plots depicting the differential expression of key genes across distinct subsets of CD4^+^ T cells. Highlighted boxes emphasize the comparative expression profiles of key genes between clonally connected SFB-specific Th17 cells in the SILP and Th1-like cells within B16-3340 tumors. One sequencing run from four T cell compartments, SILP (SFB^-^ and SFB^+^) and tumor (SFB^-^ and SFB^+^) from 5 combined mice in each group (SFB^-^ and SFB^+^).

Next, analysis of sequences of paired TCR α and β chain transcripts revealed numerous clonal relationships of CD4^+^ T cells with Th17 or follicular helper (Tfh) phenotype in the SILP and Th1-like phenotype in the B16-3340 tumors of SFB colonized mice, consistent with the intestinal origin of the tumor-infiltrating, trans-differentiated T cells that expanded in the tumor microenvironment (**Figure 3d-f**). In contrast, SFB-free mice showed minimal clonal overlap between T cells from the SILP and B16-3340 tumors, with most of these cells displaying a Th1 phenotype in the gut and a memory-like phenotype in the tumor **(Figure 3d, Extended Data Fig. 6b, c and Extended Data Table 1)**. In the SFB-colonized mice, the tumor-associated T cells sharing clonotypes with SILP T cells were characterized by upregulation of genes associated with cell trafficking, such as *Cxcr6*, chemoattraction, including *Ccl3* and *Ccl4* (potent chemoattractants for various immune cells, including cytotoxic T cells, dendritic cells, NK cells, and macrophages), pro-inflammatory cytokines *Ifng* and *Tnf*, and cytolytic functions including *Prf1*, *Klrc1,* and *Klrd1,* which together contribute to anti-tumor immunity (**Figure 3g and Extended Data Fig. 6d, e)**. In human cancers, including melanoma, breast, head and neck, and liver tumors, a comparable subset of CD4^+^ T cells with cytotoxic gene signatures was identified^32^. Recent transcriptional analysis revealed that a cytotoxic CD4^+^ T cell gene signature in bladder cancer predicts a positive response to neoadjuvant anti-PD-L1 immunotherapy^33^.

### SFB-specific T cells traffic to distal tumors

Single-cell TCR sequencing identified a clonal relationship between CD4^+^ T cells in the gut and those in the tumor microenvironment, suggesting a potential migratory pathway. To validate this observation and further investigate the connection between gut Th17 cells and tumor-infiltrating Th1-like cells in SFB-colonized mice, we employed a fate-mapping strategy. This approach allowed us to specifically track the progeny of IL-17A-expressing SFB-specific T cells that migrate from intestinal lamina propria or mesenteric lymph nodes to distal tumor sites. First, we confirmed that CD4^+^ T cells in transplanted B16-3340 tumors or the tumor-draining lymph node (TdLN) do not produce IL-17A, unlike cells from the small intestine, by using IL-17A-GFP reporter mice colonized with SFB and treated with anti-PD-1 antibody (**Extended Data Fig. 7a**). We then used IL-17A-Cre mice bred to a reporter strain (*tdTomato-ON^ΔIL-17a^* mice) to profile SFB-specific CD4^+^ T cells in the gut and implanted B16-3340 tumors in mice receiving anti-PD-1 therapy **(Figure 4a)**. In reporter mice colonized with SFB and implanted with B16-3340 tumors, more than one third of SFB Tetramer-positive and the majority of Vβ14-positive T cells within the tumors were marked as having previously expressed IL-17A. These cells were not detected either in B16-3340 tumors implanted in SFB-free mice or B16-EV tumors implanted in SFB^+^ mice (**Figure 4b, c**). As expected, tdTomato^+^Tetramer^+^ and tdTomato^+^Vβ14^+^ T cells were found in the SILP only in mice colonized with SFB (**Extended Data Fig. 7b, c**).

**Figure 4:**
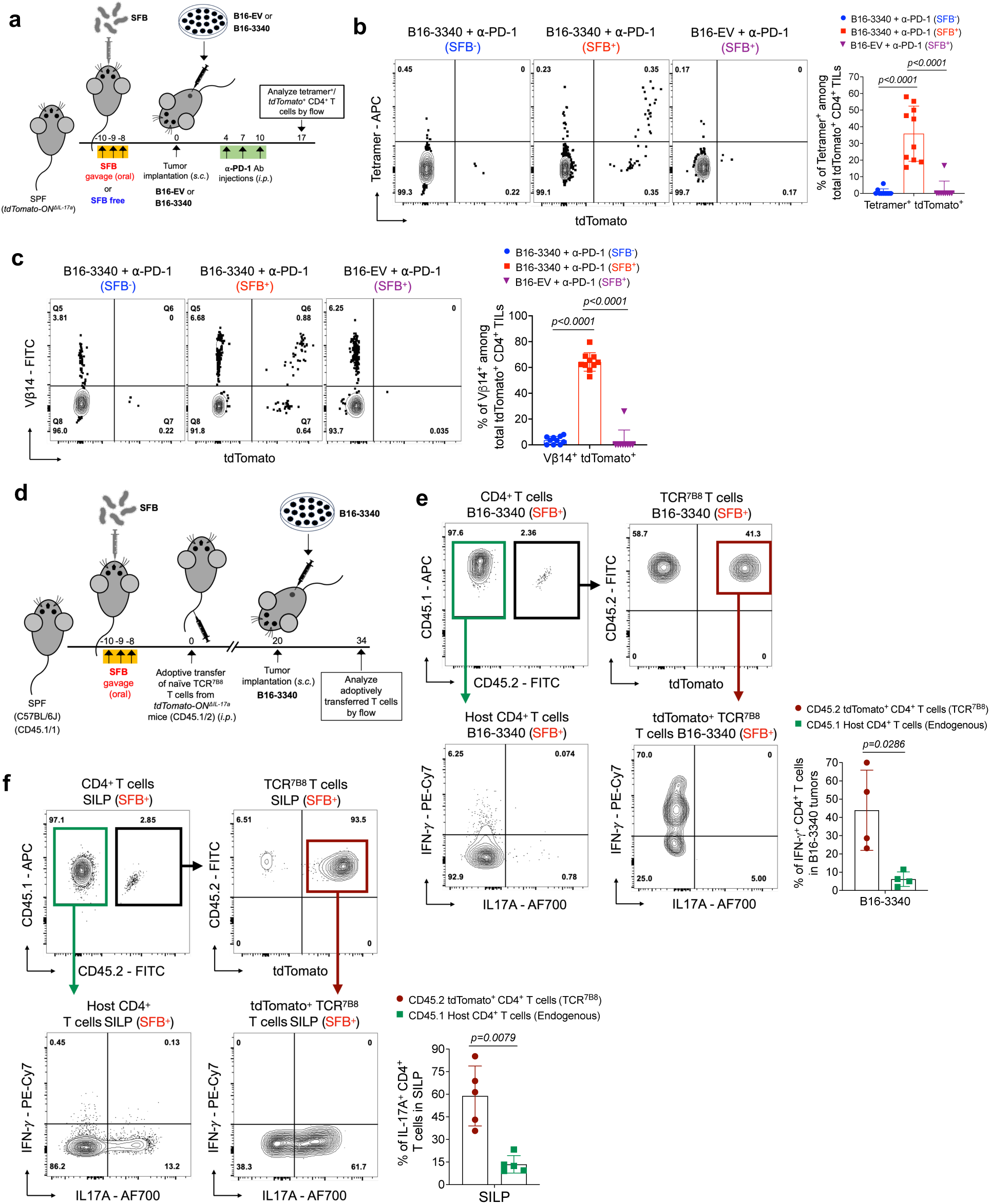
Tracking of SFB-induced T cells in gut mucosa and distal tumor tissue by MHC tetramer staining and *Il17a* fate mapping. **(a)** Schematic representation of fate mapping in *Il17a-cre;ROSALSL-tdTomato (tdTomato-ON^ΔIL-17a^)* mice, illustrating the identification of tumor-infiltrating CD4^+^ T cells that previously expressed IL-17a following colonization with SFB. (b) Ex-Th17 cell (tdTomato^+^) representation among SFB tetramer positive CD4^+^ TILs either in implanted B16-EV or B16-3340 tumors of SFB-colonized (SFB^+^) and non-colonized (SFB^-^) mice (n=10 per group). **(c)** Vβ14^+^ T cells among *tdTomato-ON^ΔIL-17a^*fate-mapped CD4^+^ cells either in implanted B16-EV or B16-3340 tumors of SFB^+^ mice compared to SFB^-^ mice (n=10 in each group). In (b) and (c), animals in both the SFB^-^ and SFB^+^ groups were administered three doses of anti-PD-1 antibody. **(d)** Schematic representation of the adoptive transfer experiment to identify tumor infiltrating SFB-specific TCR transgenic mouse T cells that previously expressed *IL17a*. Naïve T cells from *Il17a-cre;ROSALSL-tdTomato* mice bred to TCR^7B8^ transgenic mice (TCR^7B8^ *tdTomato-ON^ΔIL-17a^)* were transferred into SFB-colonized mice, and fate-mapped cells were characterized in the intestine and implanted tumors. **(e, f)** Total CD4^+^ T cells, including ex-Th17 TCR^7B8^ (tdTomato^+^) cells, were isolated from both B16-3340 tumors (e) and SILP (f) five weeks post-adoptive transfer of naïve SFB-specific TCR^7B8^ CD4^+^ T cells and activated *ex vivo* for cytokine analysis (n=5). Statistical significance shown in bar graphs was determined using unpaired two-sided Mann-Whitney t-test. Error bars denote mean ± SD. *P* values are indicated on the corresponding graphs.

We also transferred naïve CD4^+^ T cells from TCR^7B8^ *tdTomato-ON^ΔIL-17a^* transgenic mice into SFB-colonized, B6 wild-type mice. At three weeks post-adoptive transfer, B16-3340 tumors were implanted to track the migration of TCR^7B8^ Th17 T cells from the gut to distal tumor tissue (**Figure 4d**). A substantial fraction (∼50%) of the tumor-infiltrating adoptively transferred TCR^7B8^ T cells were ex-Th17 (tdTomato^+^) T cells, consistent with their migration from gut tissues (SILP and/or mLN). A large percentage of these ex-Th17 cells in the tumor tissue are significantly more proficient at producing pro-inflammatory cytokine IFN-γ compared to the endogenous tumor-infiltrating CD4^+^ T cells (**Figure 4e**). Conversely, most of the adoptively transferred T cells found in the SILP were Th17 cells (tdTomato^+^) producing IL-17A but not IFN-γ (**Figure 4f)**. The results from the above experiment suggest that SFB-specific intestinal Th17 cells migrate to distal tumors expressing SFB-3340, transforming into highly pro-inflammatory Th1-like cells.

## DISCUSSION

Molecular mimicry, with microbial antigens resembling self-antigens, has profound implications for both autoimmune disease^34^ and cancer immunotherapy. Previous studies have highlighted the potential significance of cross-reactivity between microbial antigens and tumor-associated antigens in cancers^23,35^ or autoantigens in autoimmune diseases including myocarditis, lupus, and rheumatoid arthritis^34,36,37^. Studies with cancer patients have described correlations between response to checkpoint blockade immunotherapy and the presence of specific bacterial species in the gastrointestinal tract. In a mouse model of antigen mimicry, colonization of skin with *S. epidermidis* engineered to express a model tumor antigen elicited effective T cell-mediated tumor control, consistent with the possibility that TCR cross-reactivity contributes to control of tumors in patients^24^. However, there is little understanding of how gut microbiota can be optimally enlisted to enhance immune control of distal tumors.

In this study, we aimed to establish an experimental system that would allow for mechanistic investigation of how intestinal commensal microbes could potentiate anti-tumor immunity. We took advantage of the gut-colonizing bacterium SFB because it engrafts in the presence of an intact microbiota, induces a well-characterized T cell response, and does not cause dysbiosis or inflammation, a crucial factor when investigating the influence of commensal microbes on the effectiveness of ICB. By engineering tumor cells with a synthetic SFB neoantigen, we showed that SFB-induced gut T cells could control distal tumor growth when combined with anti-PD-1 antibody treatment. The tumor-associated cells were largely derived from small intestine SFB-specific Th17 cells, and acquired Th1-like properties that were likely critical for enhancing mobilization and effector functions of tumor-infiltrating CD8^+^ T cells and other tumor-associated CD4^+^ T cells, thereby contributing to tumor control. The effector functions of the ex-Th17 cells appear superior to those of CD4^+^ T cells generated locally in the tumor-draining lymph node, potentially because of their earlier priming, resulting in effector memory function, although immunosuppressive factors in the tumor may also impair function of locally primed T cells. Moreover, ex-Th17 cells, unlike Th1 and Th17 cells, have been reported to be highly resistant to suppression by regulatory T cells, underscoring their significance in initiating a robust anti-tumor immune response^38^. Future elucidation of how microbiota-educated T cells differ from those primed in TdLN may provide valuable insights for developing more effective immunotherapies. It will be interesting to know whether gut-primed T cells can better adapt to maintain function in the tumor microenvironment than locally induced T cells that are prone to exhaustion due to continuous exposure to antigens and immunosuppressive signals and whether they may have better ability to interact with and infiltrate the tumor stroma, potentially due to specific adhesion molecules or chemokine receptors induced during gut priming.

Our single cell genomics and fate-mapping results show that intestinal bacteria-induced Th17 cells as well as Tfh cells can transition to a Th1-like state at some stage during migration into the tumor microenvironment. Different commensal bacteria can elicit distinct T cell differentiation programs, and it will be important to know whether diverse microbes differ in their capacity to induce tumor-specific T cell responses and whether differences observed can be ascribed to properties of the microbe-specific T cells. A comprehensive understanding of how different constituents of the intestinal microbiota behave in eliciting cross-reactive anti-tumor or anti-self T cell responses will likely provide opportunities for novel immune-based therapies in cancer and autoimmune disease.

## Acknowledgements

We thank members of the Littman lab for valuable discussion. We thank Susan Gottesman for valuable discussion and critical reading of the manuscript and Amanda Lund, Shruti Naik and Susan Schwab for valuable feedback. We thank the Genome Technology Center (GTC) for single cell RNA and TCR sequencing. The GTC is partially supported by NYU Cancer Center Support Grant NIH/NCI P30CA016087 at the Laura and Isaac Perlmutter Cancer Center, S10 RR023704-01A1 and NIH S10 ODO019974-01A1. This work was supported by a Merieux Foundation grant (D.R.L.), the Helen and Martin Kimmel Center for Biology and Medicine (D.R.L.), NIH grants R01AI158687 and R01CA255635 (D.R.L.), and the Howard Hughes Medical Institute (D.R.L.).

## Author Contributions

T.A.N. and D.R.L. designed the study and analyzed the data. T.A.N. performed all the experiments with assistance from G.R.M. and A.D.; T.A.N., Y.H. and Yuh.H. did bioinformatics analysis. T.A.N. and D.R.L. wrote the manuscript, with input from the other authors. D.R.L. supervised the research.

## MATERIALS AND METHODS

### Mice

Specific pathogen-free (SPF) C57BL/6J (B6) mice (both male and female, Jax #000664) were sourced from Jackson Laboratories. All transgenic mice were bred and maintained at the Alexandria Center for the Life Sciences animal facility, New York University School of Medicine, under SPF conditions. CD45.1 mice (*B6.SJL-Ptprca Pepcb/BoyJ,* Jax #002014), tdTomatoLSL mice (*B6;129S6-Gt(ROSA)26Sortm14(CAG-tdTomato)Hze/J,* Jax #007908), and IL-17a-cre mice (*Il17atm1.1(icre)Stck/J,* Jax #016879) were purchased from Jackson Laboratories. The TCR^7B8^ (7B8 TCRtg) mice were generated in our laboratory and previously described.

For IL-17A fate mapping experiments, sex-matched littermates (both males and females) were used. All mice in the experiments were 6–8 weeks of age at the onset of treatment. Animal sample sizes were determined using power analysis (power = 90%, α = 0.05) based on the mean and standard deviation from prior studies and/or pilot studies involving 4–5 animals per group. All animal procedures were conducted in accordance with protocols approved by the Institutional Animal Care and Use Committee (IACUC) of New York University School of Medicine.

### Antibodies, intracellular staining and flow cytometry

The following monoclonal antibodies were purchased from eBiosciences, BD Pharmingen or BioLegend: CD4 BUV395(GK1.5), BD 563790, 1:400; CD25 APC (PC61), Thermo Scientific 17-0251-82, 1:400; CD69 PE-Cy7 (H1.2F3), BioLegend 104512, 1:200; CD44 AF700 (IM7), BD 560567, 1:200; CD44 BV510 (IM7), BD 563114, 1:200; CD45.1 BV650(A20), BD563754, 1:400; CD45.2 FITC (104), eBioscience 11-0454-85, 1:400; CD19 PerCP-Cyanine5.5 (1D3), Tonbo Bioscience 65-0193-U100, 1:400; CD45R/B220 PerCP-Cyanine5.5 (RA3-6B2), Invitrogen 45-0452-82, 1:400; CD11c PerCP-Cyanine5.5 (N418) Invitrogen **45-0114-82**, 1:400; CD11b PerCP-Cyanine5.5 (M1/70) Invitrogen **45-0112-82**, 1:400; MHCII I-A/I-E PerCP-Cyanine5.5 (M5/114.15.2), BioLegend 107626, 1:400; NK1.1 PerCP-Cyanine5.5 (PK136), Invitrogen 45-5941-82, 1:200; TCRβ BV711 (H57-597), BD 563135, 1:200; TCRγδ PerCP-Cyanine5.5 (GL3), BioLegend 118117, 1:400; FOXP3 FITC (FJK-16s), eBioscience 11-5773-82, 1:200; RORγt BV421 (Q31-378), BD 562894, 1:200; T-BET PE-CF594 (O4–46), BD 562467, 1:70; IL-17A AF700 (TC11-18H10.1) BioLegend 506914, 1:200; IFN-γ PE-Cy7 (XMG1.2), BioLegend 505826, 1:200; Granzyme B AF700 (QA16A02), BioLegend 372222, 1:200; TNF-α BV650 (MP6-XT22), BioLegend 506333, 1:200; CD11c PE-Cy7 (N418), BioLegend 117318, 1:400; CD11b BUV395 (M1/70), BD 563553, 1:400; CXCR6 PE/dazzle 594 (SA051D1), BioLegend 151117, 1:200; CD62L PE (MEL-14) BD, Pharmingen 553151, 1:400; TCR Vβ14 FITC (14-2), BD, Pharmingen 553258, 1:400; 4′,6-diamidino-2-phenylindole (DAPI) or Live/dead fixable blue (ThermoFisher) was used to exclude dead cells.

For single-cell TCR sequencing (scTCR-seq) coupled with scRNA-seq, the following antibodies were used:

TotalSeq-C0301 anti-mouse Hashtag 1 Antibody (M1/42; 30-F11) BioLegend 155861, 1:100;

TotalSeq-C0302 anti-mouse Hashtag 2 Antibody (M1/42; 30-F11) BioLegend 155863, 1:100;

TotalSeq-C0303 anti-mouse Hashtag 3 Antibody (M1/42; 30-F11) BioLegend 155865, 1:100;

TotalSeq-C0304 anti-mouse Hashtag 4 Antibody (M1/42; 30-F11) BioLegend 155867, 1:100;

For transcription factor staining, cells were first stained for surface markers, followed by fixation and permeabilization, and then stained for nuclear factors in accordance with the manufacturer’s protocol using the FOXP3 staining buffer set from eBioscience. For cytokine analysis, cells were incubated for 3 hours in RPMI medium supplemented with 10% fetal bovine serum (FBS), phorbol 12-myristate 13-acetate (PMA) (50 ng/mL; Sigma), ionomycin (500 ng/mL; Sigma), and GolgiStop (BD Biosciences). Post-incubation, cells were stained for surface markers, fixed, permeabilized, and subsequently subjected to intracellular/nuclear cytokine and transcription factor staining according to the manufacturer’s protocol using the permeabilization buffer from eBioscience. Flow cytometric analysis was performed using an LSR II or an Aria II (BD Biosciences) and the data were analyzed with FlowJo software (Tree Star).

### Design of SFB-3340 antigen-expression construct and generation of B16-3340 and B16-EV cell lines

To develop a synthetic neoantigen mimicry model, we expressed a small, independently folded domain containing a well-characterized immunogenic CD4^+^ T cell epitope, hereafter referred to as SFB-3340, derived from a large membrane protein of Segmented Filamentous Bacterium (SFBNYU_003340, Gene Bank: EGX28318.1)^30^. A mammalian codon-optimized gene corresponding to the designed SFB-3340 antigen, linked by a flexible linker to a c-myc tag, was chemically synthesized (GenScript, USA) and cloned into the pEF1α-IRES-Neo vector (Addgene #28019) between *NheI* and *SalI* restriction sites. The expression of the antigen construct was driven by the constitutive EF-1α promoter pre-existing on this plasmid.

To establish a stable B16-F10 cell line expressing the neoantigen construct SFB-3340 (referred to as B16-3340), B16-F10 cells (ATCC #CRL-6475) were transfected with the expression plasmid complexed with TransIT-293 transfection reagent (Mirus Bio). The following day (approximately 18 to 22 hours after transfection), the B16-F10 culture media (DMEM supplemented with heat-inactivated 10% (v/v) FBS, 100 U/ml penicillin, and 0.1 mg/ml streptomycin) was replaced with selection media which is same culture media containing 1 mg/mL Neomycin (G-418) for selection. The cell cultures were then incubated for an additional 4-5 days, with the selection media changed every 2 days to select for stably transfected clones. Single clones were isolated and further expanded in selection media. The cells were passaged several times before assessing antigen expression by western blot and ex vivo activation assays. As a control cell line, B16-F10 cells were transfected with the empty vector (pEF1α-IRES-Neo without the SFB-3340 gene fragment), (referred to as B16-EV), and also selected in 1 mg/mL G-418-containing media following the same protocol as for the B16-3340 cell line.

### Immunoblotting and *ex-vivo* activation assay

To test whether B16-3340 cells are stably expressing SFB-3340 antigen and whether expressed antigen is capable of stimulating SFB-3340 antigen-specific TCR^7B8^ CD4^+^ T cells *ex-vivo*^30^, B16-3340 and B16-EV cells were lysed in M-PER reagent (Thermo Fisher Scientific) containing a protease inhibitor cocktail (Complete Mini EDTA-free; Roche) to release the antigen from the cytoplasm. The lysates were then centrifuged at 17,000g to pellet cellular debris, and supernatants were stored at -20 °C.

For western blotting, normalized amounts of proteins from the resulting cell lysates were subjected to SDS-PAGE and subsequently transferred to nitrocellulose membrane by iBlot 2 Dry Blotting System (Invitrogen). Blots were blocked in PBS blocking buffer (Licor) and then stained overnight with anti-*c-myc* antibody (1:2000, dilution, Cell signaling) at 4°C. The next day, membranes were washed three times in PBST (PBS and 0.1% Tween-20), stained with fluorescently conjugated secondary antibodies (Licor) at 1:10,000 dilution in PBS blocking buffer for 1 hour at room temperature, and then imaged in the 800-nm channel using Odyssey M Imaging system (Licor).

For the *ex-vivo* activation assay, female SPF CD45.2 mice aged 6–7 weeks were obtained from Jackson Laboratories. Spleens were harvested and processed to obtain a single-cell suspension using the GentleMACS Spleen Dissociation Kit (Miltenyi 130-095-926) following the manufacturer’s instructions. Dendritic cells (DCs) were isolated using CD11c microbeads (Miltenyi 130-125-835) as per the manufacturer’s instructions.

To isolate antigen-specific naive CD4^+^ T cells, spleens and lymph nodes from CD45.1 TCR^7B8^ transgenic mice were collected and mechanically dissociated. Red blood cells were lysed using ACK lysis buffer (Lonza). Naive TCR^7B8^ CD4^+^ T cells were sorted as CD4^+^TCRβ^+^CD44^lo^CD62L^hi^CD25^−^Vβ14^+^ using FACSAria II (BD Biosciences).

In a round-bottom 96-well plate (CELLTREAT 229190), 2×10^4^ DCs in RPMI media containing 10% FBS were incubated for 2 hours at 37°C in a 5% CO_2_ incubator with one of the following: 10 µl of PBS, 10 µl of cell lysate from 1×10^6^ B16-3340 cells, 10 µl of cell lysate from 1×10^6^ B16-EV cells, or 500 nM of chemically synthesized SFB-3340 peptide (GenScript). After the 2-hour incubation, 1×10^5^ naïve TCR^7B8^ T cells were added to each well and co-cultured for an additional 20-24 hours before staining. T cell activation was assessed by flow cytometry based on CD69 and CD25 expression.

### Colonization of mice with SFB by oral gavage

SFB colonization was achieved through three consecutive oral gavages using fecal pellets from SFB mono-associated mice, following previously described methods^30,39^. Briefly, fresh fecal pellets were homogenized through a 100-μm filter, pelleted at 3,400 rpm for 10 min, and re-suspended in PBS. Each animal was administered one-quarter pellet by oral gavage.

Colonization was confirmed by qPCR with SFB-specific primers using 16S (for fecal pellets) as housekeeping gene. Primers used were: 16S F: CGGTGAATACGTYCGG, 16S R: GGWTACCTTGTTACGACTT^40^, SFB F: GACGCTGAGGCATGAGAGCAT, SFB R: GACGGCACGGATTGTTATTCA.

### *In vivo* tumor models and antibody treatments

For B16 melanoma experiments, female SPF C57BL/6J mice aged 6-7 weeks were obtained from Jackson Laboratories and divided into two groups. The first group was maintained SFB-free, possessing SPF (SFB-minus) flora, while the second group was orally gavaged with feces from SFB mono-associated mice, as described above^30,39^. For subcutaneous tumor experiments, B16-3340 and B16-EV cells were grown in culture media with 200 μg/ml of G-418 added. Tumor cells, harvested freshly at 50-60% confluence after 3-4 passages, were resuspended in sterile PBS. Subsequently, 2.5×10^5^ cells in 100 μl PBS were injected subcutaneously into the right flank of each mouse on day 0. For experiments involving checkpoint blockade (anti-PD-1), 250 μg per mouse of anti-PD-1 (clone RMP1-14, BioXCell, BP0146) antibody was injected intraperitoneally on days 4, 7, and 10 post-tumor implantations. Tumor growth was monitored by caliper measurements, and tumor volume was calculated using the ellipsoid volume formula: 0.5 × D × d^2^, where D represents the longer diameter and d is the shorter diameter. According to our institutional animal care and use committee protocol, mice were humanely euthanized if tumors reached a volume of 2,000 mm^3^.

For *in vivo* CD4 and CD8 T cell depletion experiments, 200 μg of either anti-CD4 (clone GK1.5, BioXCell, BE0003-1) or anti-CD8a (clone 2.43, BioXCell, BE0061) antibody was injected intraperitoneally in each mouse. Injections were initiated 2 days prior to tumor implantation and continued twice weekly along with three injections of anti-PD-1 antibody (250 μg per mouse) on days 4, 7, and 10 post-tumor implantations as mentioned above^41^. The depletion efficiency was >95% in all of the mice.

### Isolation of lymphocytes from tumor, intestinal tissues and lymphoid organs

For tumor-infiltrating lymphocyte isolation, tumors were collected around days 17–18 after implantation, minced and dissociated in digestion buffer, (RPMI containing collagenase (250 U ml^−1^ of type 1 collagenase; STEMCELL technologies), DNase I (100 μg/mL; Sigma), dispase (0.1 U/ml; Worthington) and 10% FBS with constant stirring at 37 °C 30 min. After filtration, the lymphocytes were then isolated by Percoll density gradient (40%/80%) centrifugation at 800g for 20 min without brake. The interface of the Percoll layers were recovered for further analyses.

For isolation of lymphocytes from the SILP, the entire small intestine was dissected from mice. Mesenteric fat and Peyer’s patches were carefully removed from these tissues. Intestinal tissue was opened and extensively cleaned of fecal matter. This tissue was sequentially treated with HBSS 1× (1 mM DTT) at 37 °C for 10 min with gentle shaking (200 rpm), and twice with 5 mM EDTA at 37 °C for 10 min to remove epithelial cells. The remaining tissue was then minced with scissors and dissociated in RPMI containing 10% FBS, dispase (0.05 U/ml; Worthington), collagenase (1 mg/ml collagenase II; Roche) and DNase I (100 μg/ml; Sigma) with constant shaking at 37 °C for 45 min (175 rpm). The digested tissue was then filtered through a 70-μm strainer to remove large debris. Viable lamina propria lymphocytes were collected at the interface of a 40%/80% Percoll/RPMI gradient (GE Healthcare). For isolation of cells from lymph nodes and spleens, tissues were mechanically disrupted with the plunger of a 1 ml syringe and passed through 70-μm cell strainers. Red blood cells were lysed with ACK buffer (Thermo Fisher)^42^.

### MHCII tetramer production and staining

Fluorophore phycoerythrin (PE) and allophycocyanin (APC) conjugated, I-Ab/3340-A6 tetramers (SFB peptide-specific MHCII tetramers) were synthesized at the NIH tetramer core facility^43^. In brief, QFSGAVPNKT (3340-A6), an immunodominant epitope validated with the hybridoma stimulation assay, was covalently linked to I-Ab via a flexible linker, to produce pMHCII monomers. Soluble monomers were purified, biotinylated, and tetramerized with PE- or APC-labelled streptavidin^30^. Analysis of Tetramer^+^ cells was performed as previously described with minor modifications^44^. Briefly, cells were first resuspended in FACS buffer with FcR block (anti-mouse CD16/32), 2% mouse serum and 2% rat serum. Cells were then stained with PE- and APC-conjugated tetramers (10 nM) at room temperature for 1 hour in the dark.

Subsequently, the cells were washed and subjected to antibody staining against surface molecules at 4 °C.

### IFN-γ ELISPOT assay

IFN-γ ELISPOT assay was performed with a mouse IFN-γ ELISPOT kit (R&D systems) following the manufacturer’s instructions. Briefly, 5×10^4^ CD4^+^ T cells, extracted and sorted from either B16-3340 tumor tissue or SILP of SFB^+^ and SFB^-^ mice as described above. These cells were then activated with either SFB-3340 peptide (specific peptide) or Hh7-2 peptide (non-specific peptide). 2×10^4^ CD11c^+^ APC’s, purified from the spleen of SPF (SFB free) mice as described above, were used for antigen presentation in this assay. Dots were automatically enumerated with ImmunoSpot software (Version 5.0).

### Sort and adoptive transfer of naïve TCR^7B8^ (7B8 TCRtg) T cells

Recipient mice were colonized with SFB by oral gavage 7 days before adoptive transfer.

Spleens from donor TCR^7B8^ *tdTomato-ON^ΔIL-17a^* mice were harvested and mechanically disassociated. Red blood cells were lysed using ACK lysis buffer (Lonza). Naive TCR^7B8^ T cells were sorted as CD4^+^CD3^+^CD44^lo^CD62L^hi^CD25^−^TCRVβ14^+^ on the FACS Aria II (BD Biosciences). Sorted cells were then resuspended in PBS on ice and transferred into congenic isotype-labelled recipient mice by tail injection (*i.v.*). Cells from indicated tissues were analyzed 5 weeks after transfer.

### scRNA-seq and scTCR-seq experiment

For scRNA-seq experiments, the Chromium Single Cell 5′ version 2 reagent kit and Chromium Single Cell Mouse TCR Amplification Kit (10x Genomics) were used. B16-3340 tumor-infiltrated lymphocytes and lymphocytes from the SILP of SFB^-^ and SFB^+^ mice (*n=5* in each group) were extracted as described above. CD4^+^ T cells were sorted from pooled cells of either SFB^-^ or SFB^+^ tumor tissues or SILP of individual mice by FACS Aria II (BD Biosciences). Sorted CD4^+^ T cells from each group (Tumor SFB^-^, Tumor SFB^+^, SILP SFB^-^ and SILP SFB^+^) were resuspended in PBS with 0.05% BSA and stained with cell hashing antibodies, TotalSeq-C0301 to C0304 (BioLegend, Cat#155861, Cat#155863, Cat#155865, Cat#155867)^45^. After 20 minutes incubation with cell hashing antibodies, cells were washed 3 times with MACS buffer. Following cell hashing, CD4^+^ T cells from the tumor tissue of SFB^-^ and SFB^+^ groups were combined in a 1:1 ratio. Similarly, CD4^+^ T cells from the SILP of SFB^-^ and SFB^+^ groups were mixed in a 1:1 ratio. Subsequently, approximately 1.5 × 10^4^ cells per sample were loaded onto the Chromium controller (10x Genomics). Chromium Single Cell 5′ reagents were used for library preparation according to the manufacturer’s protocol. The libraries were sequenced on an Illumina HiSeq 6000. Sequencing data were aligned to the reference mouse genome mm10 with Cell Ranger (10x Genomics). The data were processed using the R packages Seurat v5^46^

### Data processing of scRNA-seq

To preprocess single-cell data, raw data from single-cell RNA sequencing (scRNA-seq) were processed using the CellRanger “multi” software (v6.1.1, 10x Genomics) with the mouse reference genome (mm10 2020-A, 10x Genomics). For single-cell T cell receptor sequencing (scTCR-seq), data were aligned and quantified using the CellRanger “multi” software (v.6.6.1, 10x Genomics) with the reference vdj_GRCm38_alts_ensembl-5.0.0, using default settings.

For scRNA-seq analysis, we excluded cells with fewer than 200 detected genes. Cells manifesting more than 5% mitochondrial genes were also removed from the dataset. HTO counts were normalized using centered log ratio transformation before hashed samples were demultiplexing using the Seurat::HTODemux function (positive quantile set to 0.99). Doublets mapped to multiple HTO tags were removed. RNA counts were normalized using Seurat::SCTransform function with regressions of cell cycle score, ribosomal and mitochondrial percentages^47^. Sample integrations were performed for paired SFB^-^ and SFB^+^ samples from the same tissue using Seurat standard scRNSeq integration workflow with 3000 anchor genes. A shared nearest neighbor graph was then built based on the first 40 principal components (PCs) followed by Leiden clustering using Seurat::FindClusters function at multiple resolutions in order to identify potential rare cell types^48^. Cluster phenotyping was based on canonical cell type markers and differentially expressed genes between clusters identified using Seurat::FindAllMarkers function with a logistic regression model. Cells were then projected onto a Uniform Manifold Approximation and Projection (UMAP) for visualization^49^.

TCR sequence data were processed using Cell Ranger vdj pipeline to identify TCR genes and CDR3 sequences. For each sample, full length, productive TRB and TRA chains were retained for downstream analysis. Clonal expansions were determined using nucleotide sequences of CDR3 sequence of both chains that appeared in at least 3 cells from all samples. TCR information was added as metadata into the scRNASeq Seurat object based on cell barcodes and sample information. To characterize T cell phenotypes of TCR clonotypes, a metadata file was generated from the Seurat object and analyzed and quantified using Microsoft Excel (v.16.73).

Differential expression of genes between two groups was tested with MAST package (MAST version by Seurat v5 is MAST_1.28.0), which uses a hurdle model tailored to scRNA-seq data^50^. Genes with an adjusted *p*-value smaller than 0.05 (based on Bonferroni correction using all genes in the dataset) were considered as statistically significant.

### Statistical analysis

Unpaired two-sided *t*-test, paired two-sided *t*-test, one-way ANOVA with multiple comparisons with Bonferroni correction, two-way ANOVA with multiple comparisons and Bonferroni correction, Mann–Whitney test and the Mantel–Cox test (for survival curves) were performed to compare the results using GraphPad Prism version 9 (GraphPad). No samples were excluded from analysis. Exact *P* values are provided where possible. *P* < 0.05 are considered as significant.

---

**Extended Data Fig. 1:**
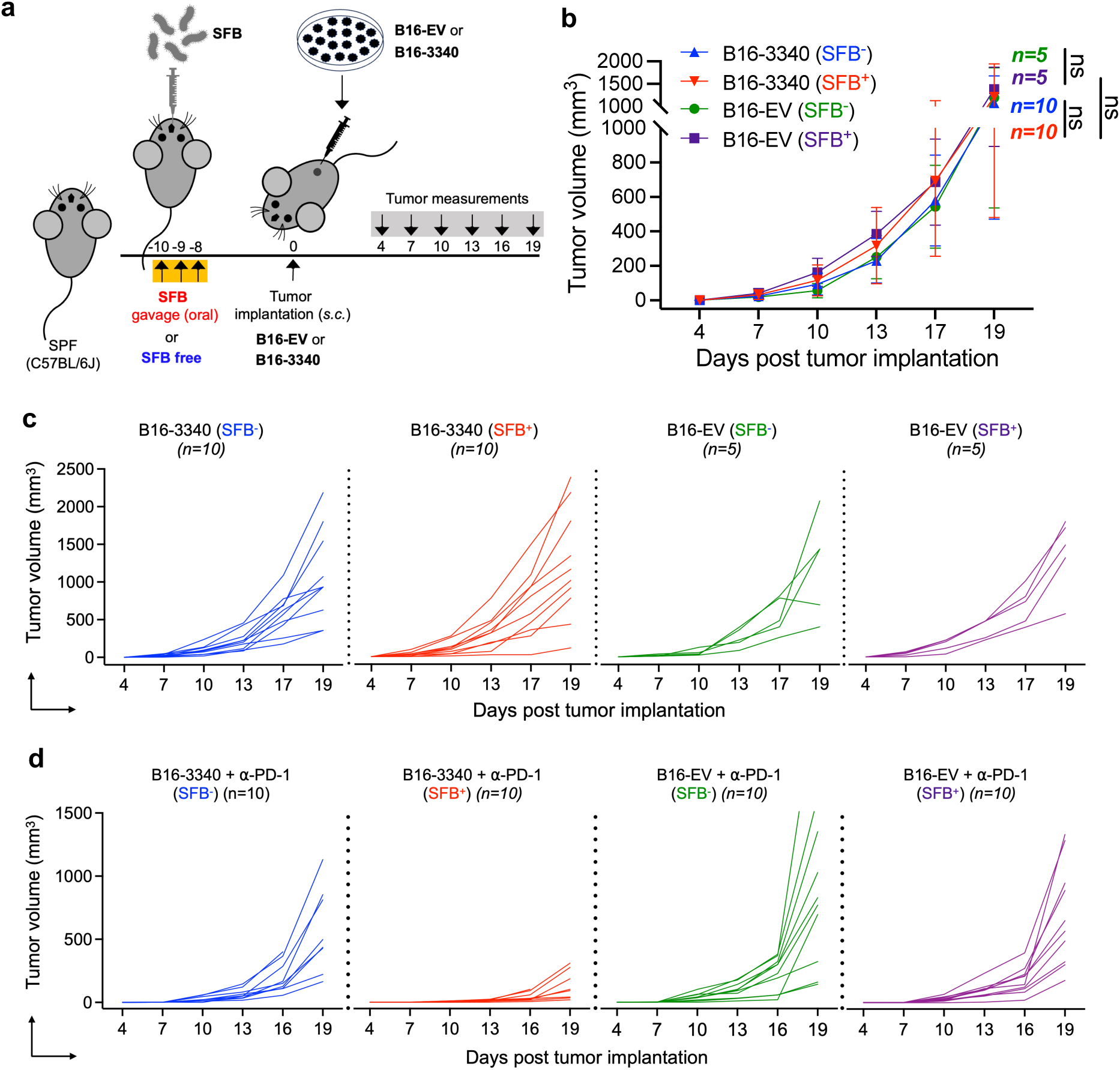
SFB colonization alone (without anti-PD-1 treatment) does not show significant change in tumor growth. **(a)** Experimental strategy for testing the artificial mimicry model in C57BL/J mice without anti-PD-1 antibody treatment. Mice were colonized with SFB or kept SFB-free followed by tumor implantation either with B16-3340 or B16-EV tumor cells. No anti-PD-1 treatment was administered. **(b)** Caliper measurements are shown as growth curves of B16-3340 (n=10) or B16-EV (n=5) implanted tumors. Without ant-PD-1 antibody treatment, no significantly change in tumor growth was observed between SFB^-^ and SFB^+^ mice. (**c)** Tumor volume growth curves over time, each line representing the tumor size of an individual mouse in the group, blue: B16-3340 (SFB^-^), red: B16-3340 (SFB^+^), green: B16-EV (SFB^-^), and magenta: B16-EV (SFB^+^). **(d)** Related to Figure 1 **(e)**, tumor growth curves over time in individual mice in SFB^+^ and SFB^-^ groups that were treated with anti-PD-1 antibody. Each line represents the tumor size of an individual mouse in the group, blue: B16-3340 + α-PD-1 (SFB^-^), red: B16-3340 + α-PD-1 (SFB^+^), green: B16-EV + α-PD-1 (SFB^-^), and magenta: B16-EV + α-PD-1 (SFB^+^).

**Extended Data Fig. 2:**
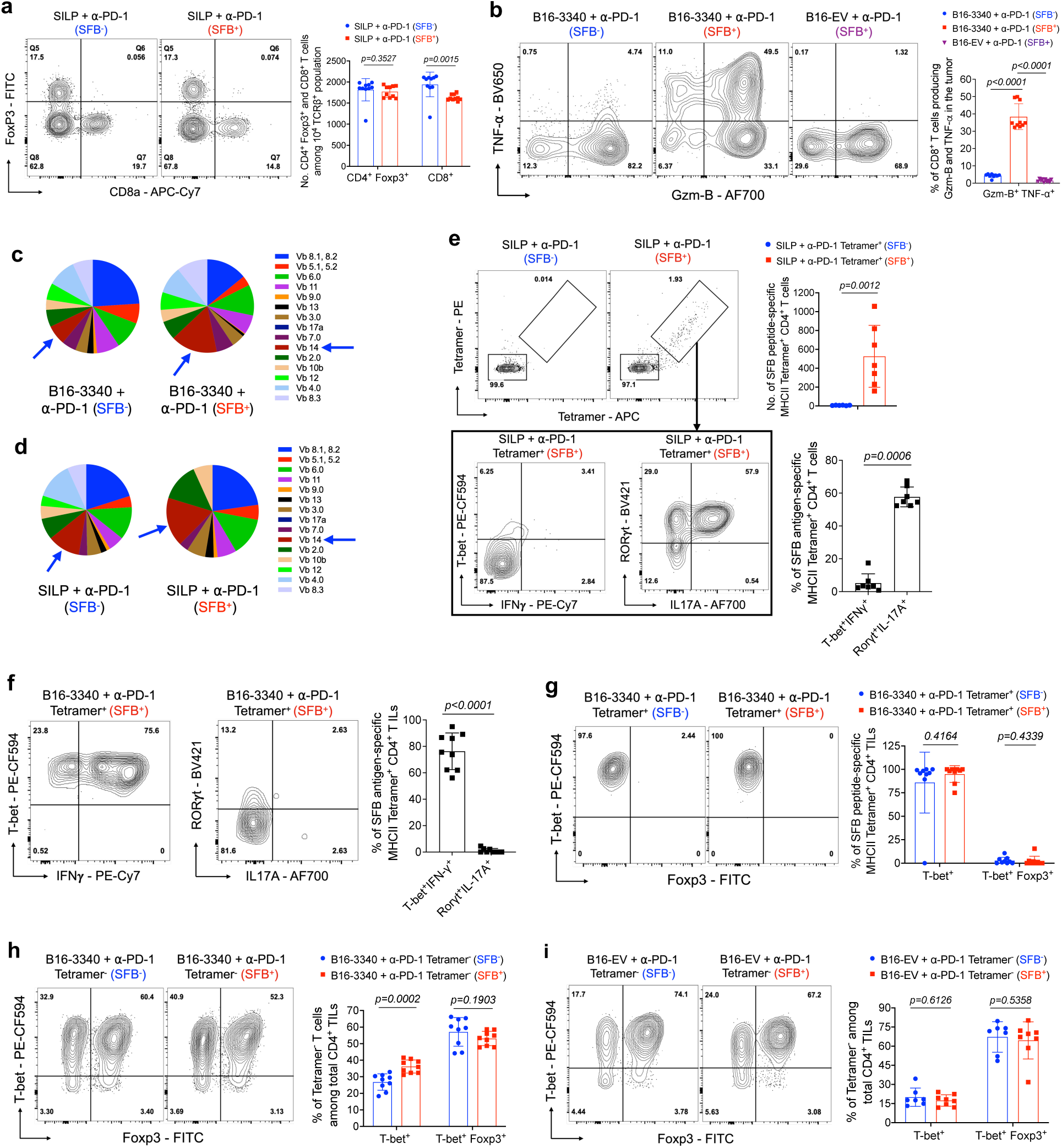
Differential immune responses in the SILP and tumor microenvironment of SFB-colonized mice after anti-PD-1 therapy. **(a)** Representative flow cytometry plots of CD8^+^ T cells and regulatory T cells (Tregs; CD4^+^ Foxp3^+^) in the SILP of SFB-free (SFB^-^) and SFB-colonized (SFB^+^) mice (left panel) and quantification of the frequencies of CD8^+^ T cells and Tregs in the SILP across these groups (right panel, n=10 per group). **(b)** Representative flow cytometry plots showing expression of effector gene products (Gzm-B^+^ and TNF-α^+^) in CD8^+^ TILs isolated from B16-3340 tumors in SFB^-^ and SFB^+^ mice and B16-EV tumors in SFB^+^ mice (left). The right panel quantifies the expression levels of these markers across the different groups (n=9 per group for B16-3340 and n=6 for B16-EV tumors). (**c**) Representation of TCR Vβ usage among CD4^+^ T cells in B16-3340 tumor tissue of SFB^-^ and SFB^+^ mice, respectively. (**d**) Analysis of TCR Vβ repertoire usage among CD4^+^ T cells in SILP of SFB^-^ and SFB^+^ mice. **(e)** Top, SFB peptide-specific MHCII tetramer staining of CD4^+^ T cells isolated from the SILP of SFB^-^ and SFB^+^ mice, with quantification of results shown on the left (bar diagram, n=7). Bottom, expression of IFN-γ and T-bet, as well as IL-17A and RORγt in Tetramer^+^ CD4^+^ T cells from the SILP of SFB^+^ mice, with corresponding quantification shown on the right (n=7). (**f**) Expression of IFN-γ and T-bet, as well as IL-17A and RORγt in Tetramer^+^ CD4^+^ T cells isolated from the B16-3340 tumors of SFB^+^ mice, with corresponding quantification shown on the right (n=9). (**g**) Foxp3 and T-bet expression profiles of Tetramer^+^ CD4^+^ TILs isolated from the B16-3340 tumors of SFB^-^ and SFB^+^ mice with corresponding quantification shown on the right (n=9 per group). (**h, i**) Foxp3 and T-bet expression profiles of Tetramer^-^ CD4^+^ TILs isolated from the B16-3340 tumors (h) and B16-EV tumors (i) of SFB^-^ and SFB^+^ mice (n=8 to 10 per group). The corresponding bar graphs quantify the expression levels of these transcription factors across the different groups. In each experiment, three doses of anti-PD-1 antibody were administered to the animals in both the SFB^-^ and SFB^+^ groups. For cytokine expression analysis, cells were activated *ex vivo* by PMA/Ionomycin for 3 hours at 37°C. Statistical significance was determined using unpaired two-sided Mann-Whitney t-test. Error bars denote mean ± SD. *P* values are indicated on the corresponding bar graphs.

**Extended Data Fig. 3:**
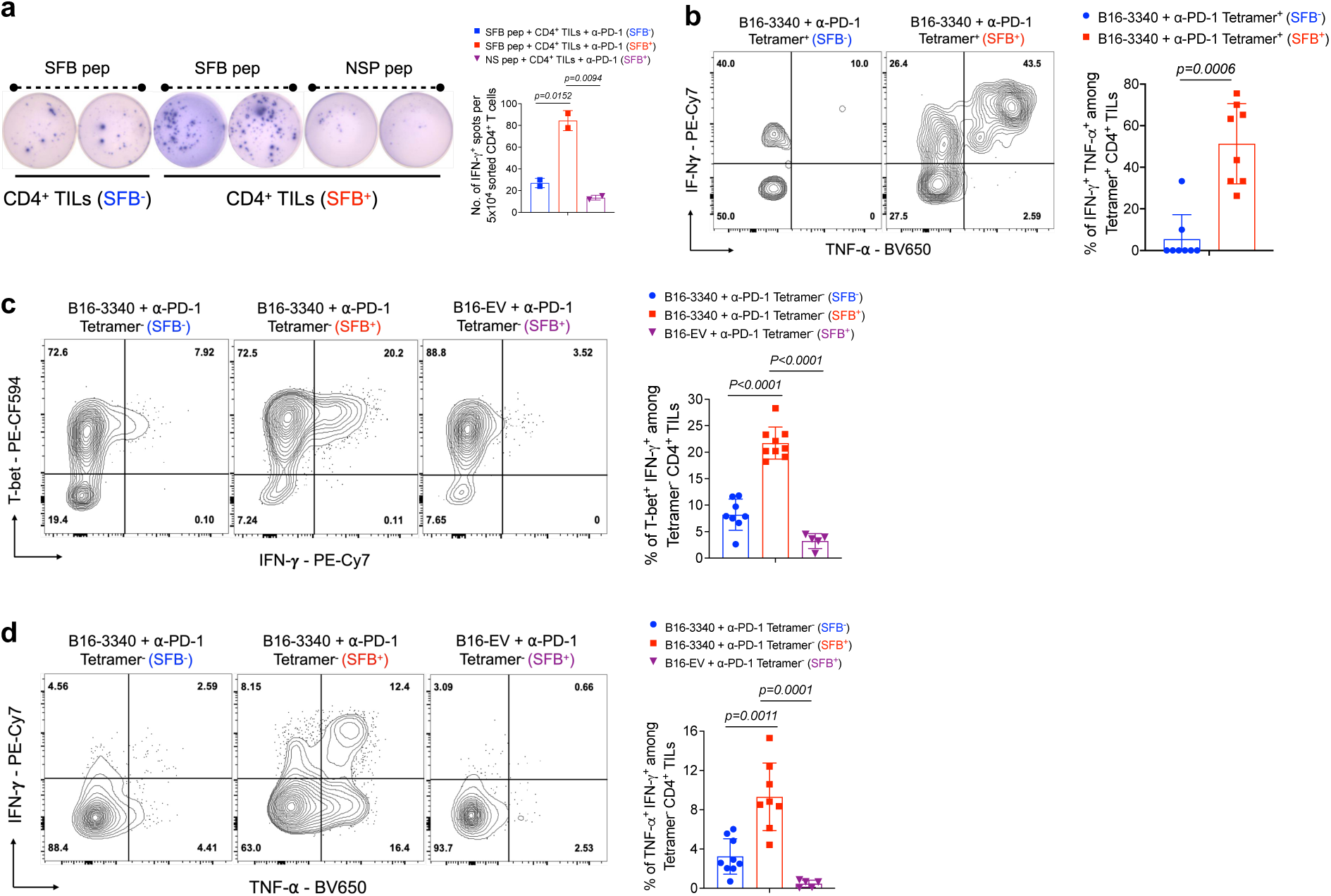
Effects of SFB colonization on immune responses in B16-3340 tumors and SILP after anti-PD-1 antibody treatment. **(a)** IFN-γ ELISpot assay of CD4^+^ TILs isolated from B16-3340 tumors in SFB^-^ and SFB^+^ mice and activated for 24h with SFB peptide (SFB peptide recognized by TCR^7B8^) or non-specific peptide (NSP) *ex vivo.* (**b**) Representative flow cytometry plots depicting the expression of effector gene products, TNF-α and IFN-γ, in Tetramer^+^ CD4^+^ TILs isolated from B16-3340 tumors in SFB^-^ and SFB^+^ mice (left) and quantification of cytokine expression across the two groups (right, n=8 per group). **(b)** Representative flow cytometry plots showing the expression of transcription factor T-bet and cytokine (TNF-α and IFN-γ) in Tetramer^-^ CD4^+^ T cells isolated from B16-3340 tumors of SFB^-^ and SFB^+^ mice (n=8 to 9 per group) as well as from B16-EV tumors in SFB^+^ mice (n=5).The corresponding bar graphs, associated with each flow cytometry plot, quantifies the expression levels of transcription factors and cytokines across the different groups. In all experiments, mice in both SFB^-^ and SFB^+^ groups received three doses of anti-PD-1 antibody. For cytokine expression analysis, cells were activated *ex vivo* by PMA/Ionomycin for 3 hours at 37°C. Statistical significance was determined using unpaired two-sided Mann-Whitney t-test. Error bars denote mean ± SD. *P* values are indicated on the respective graphs.

**Extended Data Fig. 4:**
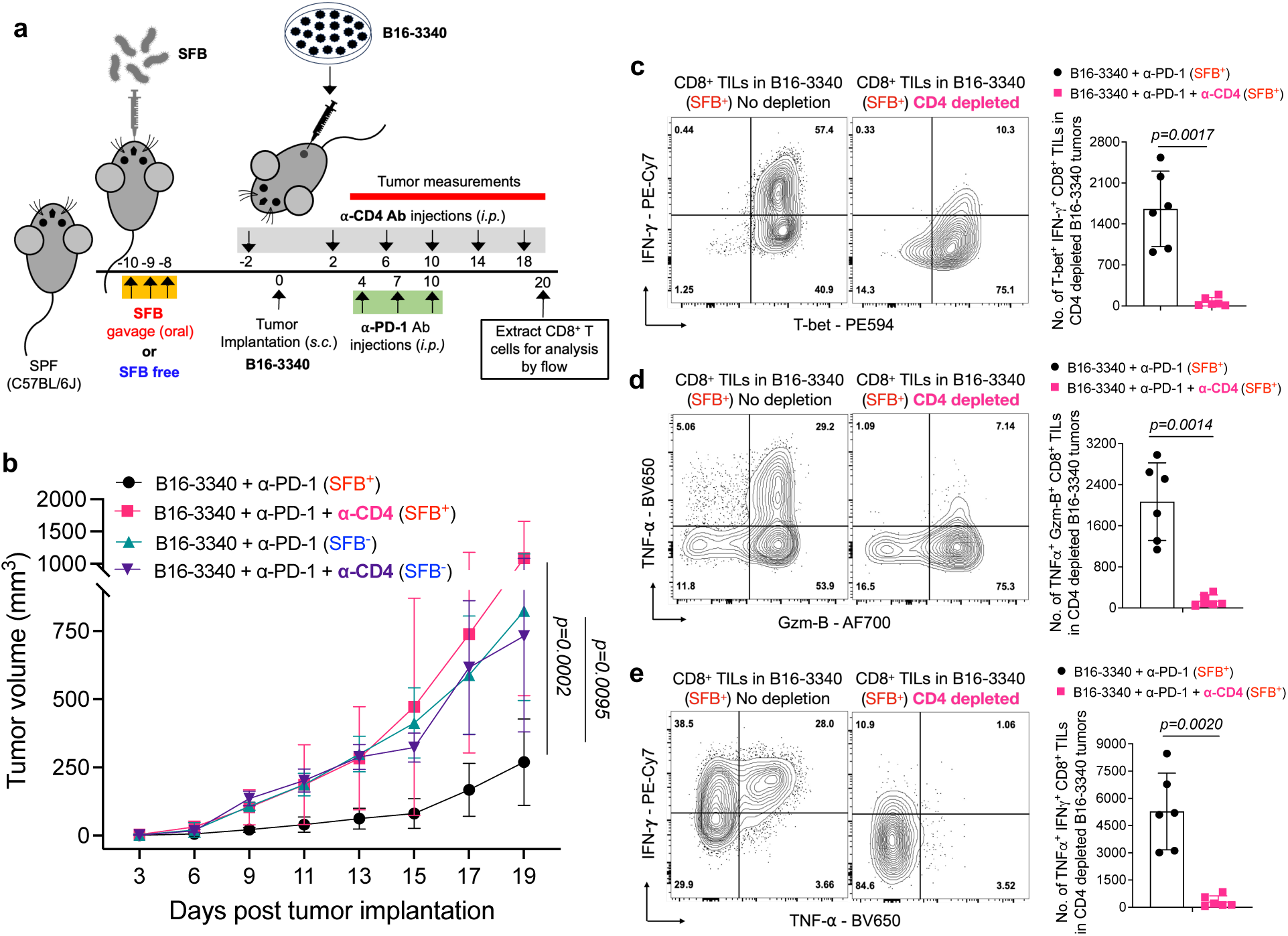
Effects of *in vivo* CD4 T cell depletion on tumor growth and CD8^+^ TIL effector function in SFB^+^ and SFB^-^ mice treated with anti-PD-1 antibody. **(a)** Schematic representation of *in-vivo* CD4 T cell depletion in SFB^+^ and SFB^-^ mice with B16-3340 tumors treated with anti-PD-1 antibody. All mice received 3 injections of anti-PD-1 Ab (250 µg/mouse *i.p.* on days 4, 7 and 10 post tumor implantation) with or without *in vivo*-depleting anti-CD4 mAb twice per week (200µg/mouse *i.p.,* on days -2, 2, 6, 10, 14 and 18 post-tumor implantation) (n=6 per group). **(b)** Caliper measurements are shown as growth curves of B16-3340 implanted tumors in SFB^+^ and SFB^-^ mice, with or without *in vivo*-depletion of CD4 T cells using anti-CD4 monoclonal antibody (n=6 per group). Statistical significance was calculated using two-way ANOVA and Sidak’s multiple comparisons. **(c-e)** Expression of effector gene products (IFN-γ, TNF-α and Gzm-B) in CD8^+^ TILs from B16-3340 tumors in SFB colonized mice treated with anti-PD-1 antibody, with and without *in vivo*-depletion of CD4^+^ T cells (n=6 per group). For cytokine expression analysis, cells were activated *ex vivo* by PMA/Ionomycin for 3 hours at 37°C. Statistical significance was determined using unpaired two-sided Mann-Whitney t-test. Error bars denote mean ± SD. *P* values are indicated on the corresponding graphs.

**Extended Data Fig. 5:**
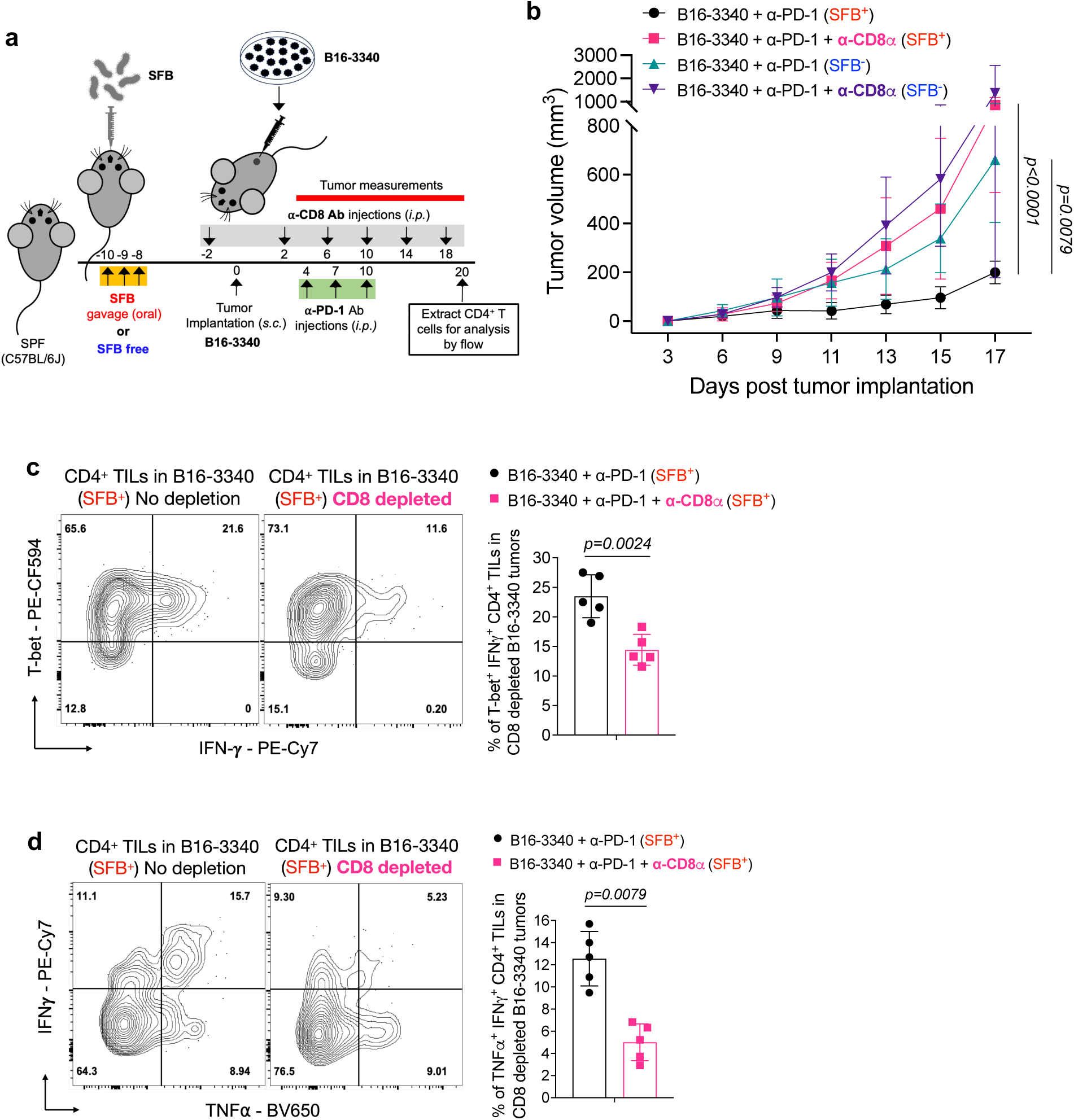
Effect of *in-vivo* CD8 T cell depletion on anti-PD-1 therapy efficacy in SFB^+^ and SFB^-^ mice with B16-3340 tumors. **(a)** Schematic representation of *in-vivo* CD8 T cell depletion in SFB^+^ and SFB^-^ mice with B16-3340 tumors treated with anti-PD-1 antibody. All mice received 3 injections of anti-PD-1 Ab (250 µg/mouse *i.p.* on days 4, 7 and 10 post tumor implantation) with or without *in vivo*-depleting anti-CD8 mAb twice per week (200µg/mouse *i.p.,* on days -2, 2, 6, 10, 14 and 18 post-tumor implantation) (n=5 for each group). **(b)** Caliper measurements are shown as growth curves of B16-3340 implanted tumors in SFB^-^ and SFB^+^ mice treated with anti-PD-1, with or without *in vivo*-depleting anti-CD8 monoclonal antibody. **(c, d)** Expression of effector gene products (IFN-γ and TNF-α) in CD4^+^ TILs from B16-3340 tumors of SFB colonized mice treated with anti-PD-1 antibody, with and without administration of *in vivo*-depleting anti-CD8 antibody (n=5 per group). For cytokine expression analysis, cells were activated *ex vivo* by PMA/Ionomycin for 3 hours at 37°C. Statistical significance, shown on the survival graph is calculated using two-way ANOVA and Sidak’s multiple comparisons and statistical significance, shown on the bar graphs, was determined using unpaired two-sided Mann-Whitney t-test. Error bars denote mean ± SD. *P* values are indicated on the corresponding graphs.

**Extended Data Fig. 6:**
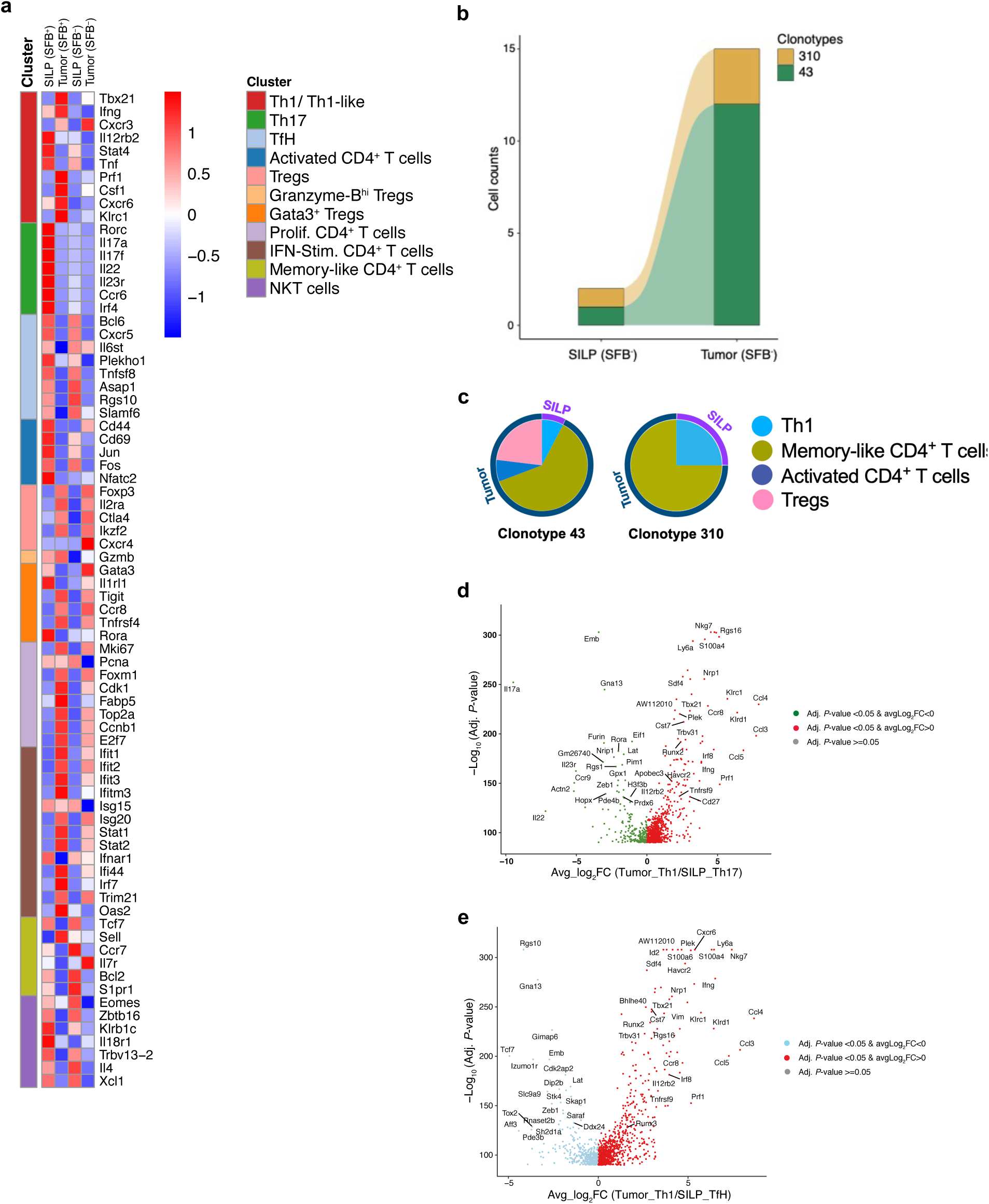
Gene expression in CD4^+^ T cells within SILP and tumors. **(a)** Heatmap showing normalized gene expression scaled by row (gene) of top differentially expressed and key cell lineage marker genes. (**b**) An alluvial graph depicting the clonal connectivity of CD4^+^ T cell clonotypes between the SILP and tumor tissues in SFB-free mice. Each block in the bar diagram represents cell counts within a distinct CD4^+^ T cell clonotype, with branches in the graph illustrating the shared clonotypes between SILP and tumor compartments. (**c**) Clonal expansion with phenotypic switching (represented by color) within the tumor for two shared clonotypes originating from the gut in SFB-free mice. **(d, e)** Volcano plots highlighting gene expression differences between Th17 (b) and TfH (c) cells in the SILP and Th1 cells in the tumor. Statistical significance was determined using the MAST package, with color coding indicating the magnitude of change: Red for upregulated genes and green or light-blue for downregulated genes (only significant genes are shown, genes with adjust-p-value >=0.05 are not included in the plot).

**Extended Data Fig. 7:**
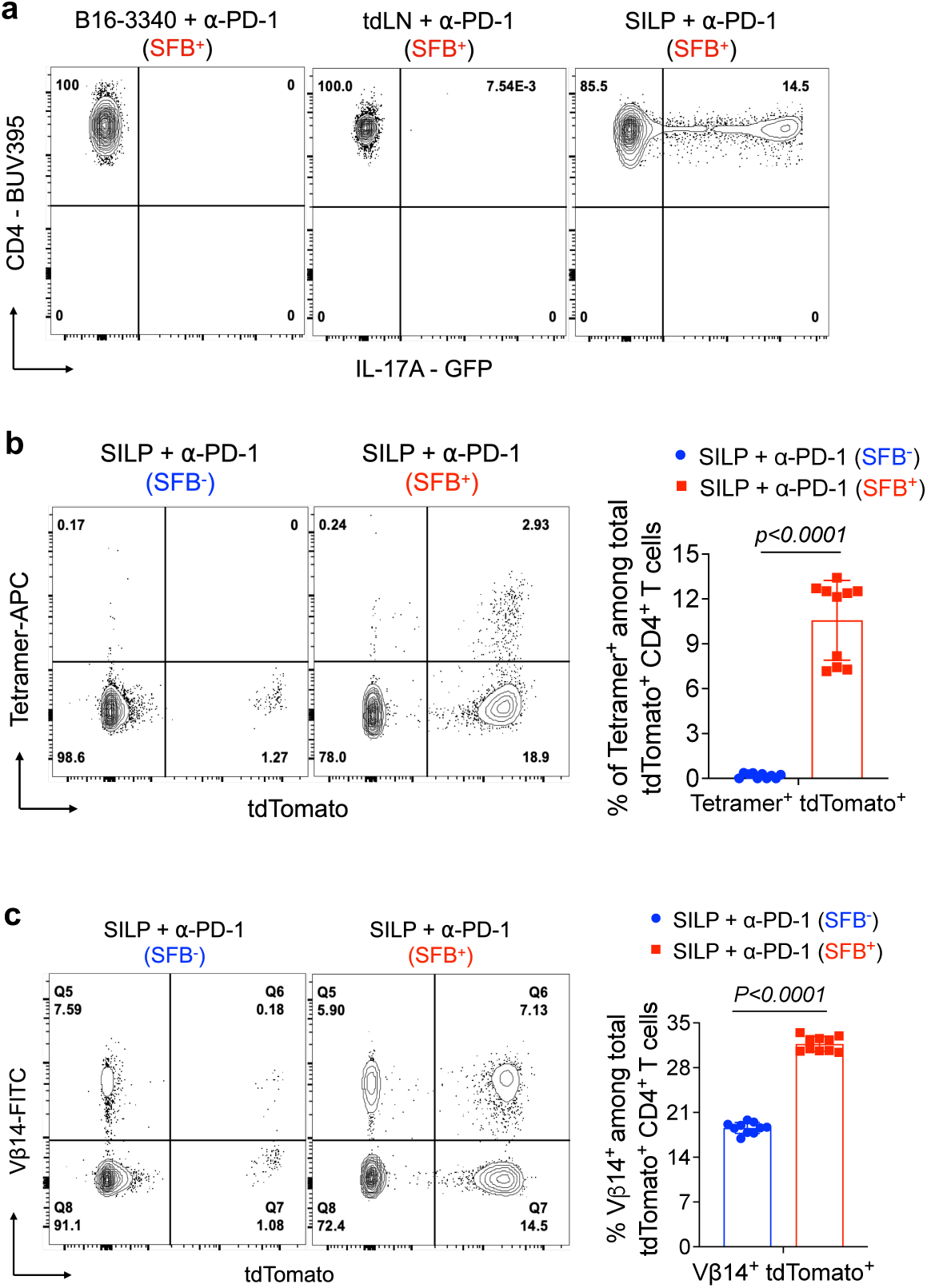
Characterization of Il17a fate mapped T cells in the small intestine of SFB-colonized mice. **(a)** GFP expression in CD4^+^ T cells isolated from B16-3340 tumor tissue, tumor draining lymph node (TdLN) and SILP of IL-17A-GFP reporter mice colonized with SFB and treated with anti-PD-1 antibody. **(b)** Current or previous *Il17a* expression (tdTomato^+^) among SFB tetramer positive CD4^+^ T cells in the SILP of *tdTomato-ON^ΔIL-17a^*fate-mapped mice with and without SFB colonization (n=10 per group). **(c)** Vβ14^+^ T cells among *tdTomato-ON^ΔIL-17a^*fate-mapped CD4^+^ cells in SILP of SFB^+^ mice compared to SFB^-^ mice (n=10 in each group). In (b) and (c), mice in both the SFB^-^ and SFB^+^ groups received three doses of anti-PD-1 antibody. Statistical significance shown in (b) and (c) was determined using unpaired two-sided Mann-Whitney t-test. Error bars denote mean ± SD. *P* values are indicated in the corresponding graphs.

**Extended Data Fig. 8:**
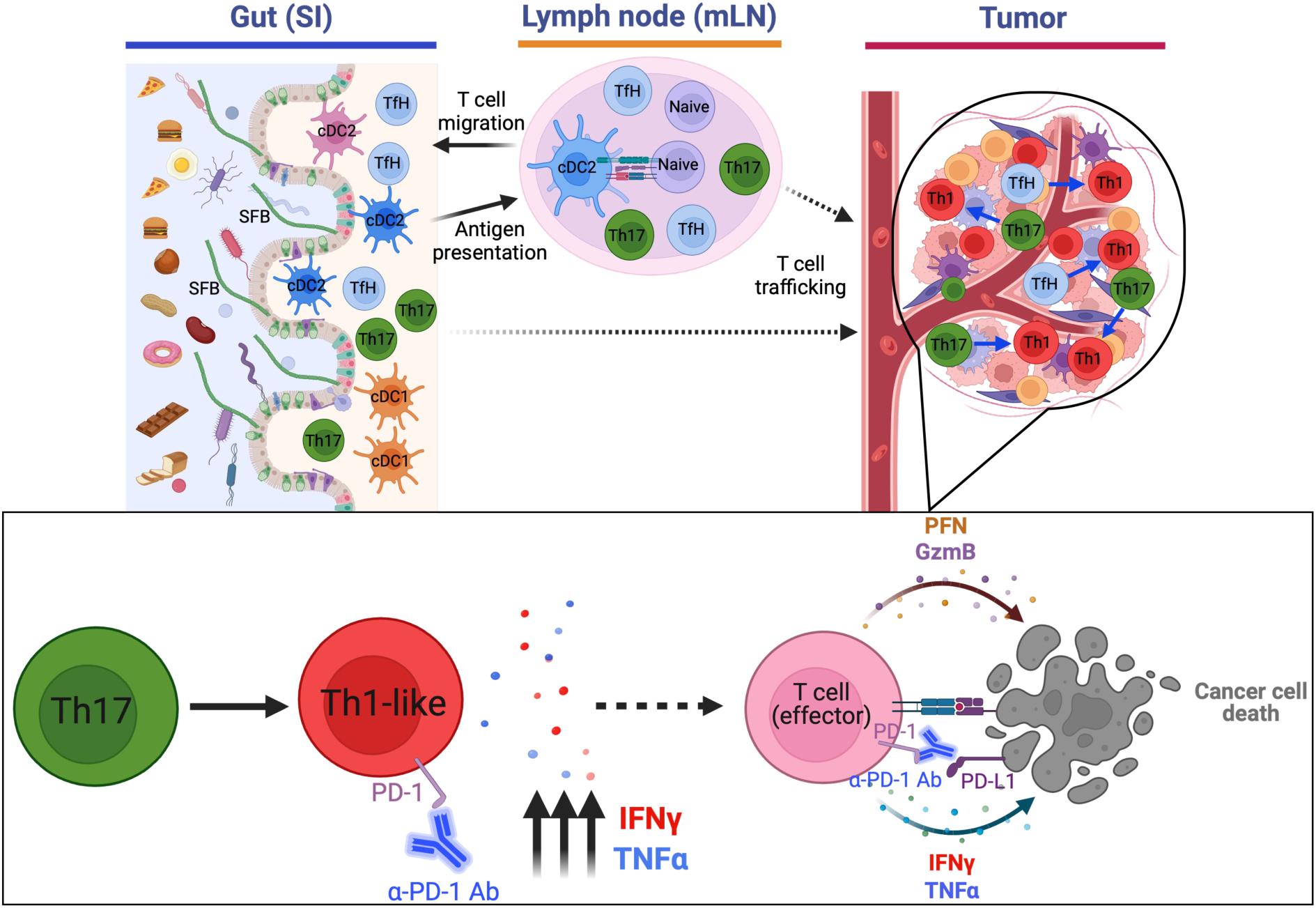
Schematic illustration of how gut commensal-induced T cells affect cancer immunotherapy in a synthetic mimicry model. SFB colonization induces antigen-specific CD4^+^, and likely CD8^+^ T cells, which then distribute to other parts of the body but are retained and subsequently expand only in the tumor tissue which expresses SFB antigen (B16-3340 tumors). Th17 cells specific for SFB-3340 antigen transdifferentiate into Th1 cells, likely in the tumor tissue under the influence of the tumor microenvironment, and produce proinflammatory cytokines IFN-γ and TNF-α, which aid in the infiltration and effector capabilities of cytotoxic T cells specific for the tumor.

